# A *C. elegans* male pheromone feminizes germline gene expression in hermaphrodites and imposes life-history costs

**DOI:** 10.1101/2023.02.17.528976

**Authors:** David Angeles-Albores, Erin Z. Aprison, Svetlana Dzitoyeva, Ilya Ruvinsky

## Abstract

Sex pheromones improve reproductive success, but also impose costs. Here we show that even brief exposure to physiological amounts of the dominant *C. elegans* male pheromone, ascr#10, alters the expression of thousands of genes in hermaphrodites. The most dramatic effect on the transcriptome was the upregulation of genes expressed during oogenesis and downregulation of genes associated with male gametogenesis. Among the detrimental effects of ascr#10 on hermaphrodites is the increased risk of persistent infections caused by pathological pharyngeal hypertrophy. Our results reveal a way in which social signals help to resolve the inherent conflict between spermatogenesis and oogenesis in a simultaneous hermaphrodite, presumably to optimally align reproductive function to the presence of potential mating partners. They also show that the beneficial effects of the pheromone are accompanied by harmful consequences that reduce lifespan.

## Introduction

Sex pheromones alter multiple aspects of the recipient’s reproductive biology (Wyatt 2014). We are studying these phenomena in a nematode *Caenorhabditis elegans*. In this species, hermaphrodites typically reproduce by selfing, but retain the ability to mate with males that are rare in both wild and laboratory populations (Frezal and Felix 2015). As is common in other species, “the *C. elegans* pheromone” is a complex blend of molecules (Srinivasan, et al. 2008; Srinivasan, et al. 2012), including compounds that are predominantly produced by males (Ludewig, et al. 2019; Burkhardt, et al. 2023). The best studied male pheromone, the ascaroside ascr#10 (Izrayelit, et al. 2012), at low physiological concentrations (Aprison and Ruvinsky 2017) can modulate several reproduction-related traits in hermaphrodites. For example, it improves oocyte quality (Aprison, et al. 2022) and increases stores of germline precursors (Aprison and Ruvinsky 2016).

The specific molecular mechanisms by which ascr#10 alters hermaphrodite physiology remain largely unknown. It is reasonable to posit, however, that the reproductive system is a major target of this pheromone. *C. elegans* is a simultaneous hermaphrodite. Morphologically, hermaphrodites resemble females, but they make a cache of sperm prior to irreversibly switching to oocyte production (Kimble and Crittenden 2007). Because hermaphrodites have reproductive organs of both sexes, they must balance resource investment between male and female functions that are inherently in conflict (Charnov 1979; Parker 2006). To males, the value of hermaphrodites resides in oocytes and other “female” traits, whereas hermaphrodite sperm and other “male” traits are a source of potential competition. We therefore hypothesized that one likely effect of the male pheromone is to bias hermaphrodites toward feminization, at the expense of male functions (Figure 1a).

**Figure 1.**
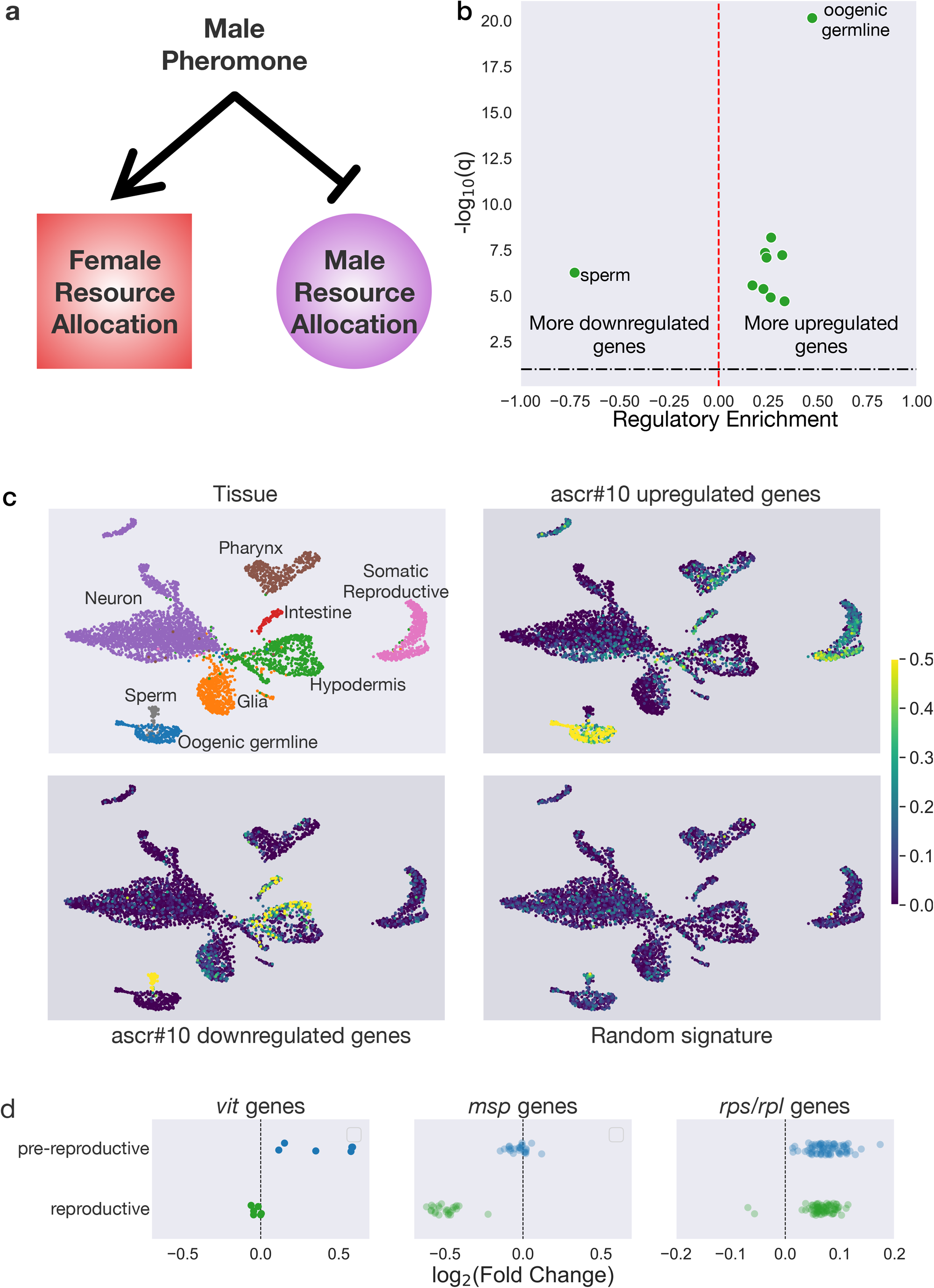
Transcriptomic signature of shifting sex allocation in the presence of ascr#10. **a**, The hypothesis tested in this study – exposure of hermaphrodites to the male pheromone is expected to increase allocation to “female” reproductive functions and decrease allocation to “male” reproductive functions. **b**, Regulatory enrichment scores of tissues in response to ascr#10 in reproductive animals. A positive regulatory enrichment score indicates that the fraction of genes that were differentially up-regulated in response to ascr#10 in that tissue is greater than expected by random chance. Only significantly enriched tissues are shown. **c**, UMAP plot of single-cell RNA-seq data showing the tissue clusters. ascr#10 upregulated genes are enriched in the oogenic germline, the somatic reproductive tissues, and the pharynx. Downregulated genes are expressed predominantly in hermaphroditic sperm, and the hypodermis. A random signature is shown for comparison. **d**, Coherent differential expression of genes encoding vitellogenins (*vit*), Major Sperm Protein (*msp*), and ribosomal protein subunits (*rpl, rps*) implies that these groups of genes behave as functional units.

## Results and discussion

### ascr#10 feminizes the hermaphrodite transcriptome by altering expression of thousands of genes

Because altered gene expression likely undergirds physiological effects of the pheromone, we assessed changes caused by ascr#10 in the global transcriptional profile of hermaphrodites (Aprison, et al. 2022). Pheromone exposures were brief and took place after the switch from spermatogenesis to oogenesis. We assayed two timepoints – a few hours before and a few hours after the onset of egg-laying; we refer to these as pre-reproduction and reproduction, respectively.

We measured 12,244 genes across both timepoints. We identified 1,663 differentially expressed genes in pre-reproductive animals and 3,622 genes in reproductive animals in response to ascr#10. The magnitude of differential expression was modest (<2 fold for almost all genes), but changes were positively correlated across timepoints (Supplementary Information). Aggregating weakly differentially expressed genes (those that show modest fold change) can be used to identify changes in organismal state (Angeles-Albores, et al. 2017) or to reconstruct genetic pathways (Angeles-Albores, et al. 2018).

Using previously curated WormBase annotations that assigned genes to tissues in which they are expressed, we evaluated whether differentially expressed genes were biased to specific cell types. Although upregulated genes were distributed across several tissues, the most notably overrepresented category was the oogenic germline, while the only downregulated tissue was sperm (Figure 1b).

A potential downside to this analysis is that it represents expression data as binary values (positive or negative) and relies on albeit expert, nonetheless human annotations. To overcome these limitations, we verified our results using a single-cell whole organism RNA-seq dataset (Taylor, et al. 2021). We derived stringent signatures of genes up- or downregulated by ascr#10 and computed expression-normalized scores per signature. We compared these scores to a random signature for reference. In broad agreement with the analysis based on curated annotations (Figure 1b), we saw that the ascr#10 upregulated signature was enriched in the oogenic germline and the female reproductive soma, whereas the ascr#10 downregulated signature was enriched among the sperm-expressed genes (Figure 1c).

Changes in expression in three functional classes of genes help to illustrate the response of the hermaphrodite transcriptome to ascr#10 (Figure 1d). Yolk accounts for over 1/3 of total protein in the embryo, while vitellogenins that encode yolk proteins constitute ∼3% of total adult mRNA (Perez and Lehner 2019). Expression of all vitellogenins was increased in pre-reproductive adults. The lack of enrichment in reproductive adults may be due to production being saturated. In reproductive adults, Major Sperm Protein (*msp*) genes (Miller, et al. 2001) were downregulated. Finally, all genes encoding ribosomal proteins were upregulated, indicating increased demand for translational machinery. This result is consistent with the idea of greater offspring provision and the prior observation of increased ribosomal gene expression in mated hermaphrodites (Booth, et al. 2019).

### ascr#10 increases pharyngeal pumping and causes pharyngeal hypertrophy in hermaphrodites

Because ascr#10 increases expression of pharyngeal genes (Figure 1b, c), we quantified the effects of this male pheromone on pharyngeal function and observed significantly increased pumping rate (Figure 2a). We hypothesized that chronically increased pumping could lead to pharyngeal hypertrophy. Indeed, hermaphrodites raised in the presence of ascr#10 had a significantly enlarged posterior bulb of the pharynx (Figure 2b). A similar swollen pharynx phenotype is a prevalent life-limiting pathology identified in a necropsy analysis, with associated deaths proximally caused by bacterial infection (Zhao, et al. 2017). Hermaphrodites aging in the presence of ascr#10 showed considerably increased presence of live bacteria in the intestine (Figure 2c), suggesting that the pheromone-induced pharyngeal hypertrophy likely contributes to accelerated senescence.

**Figure 2.**
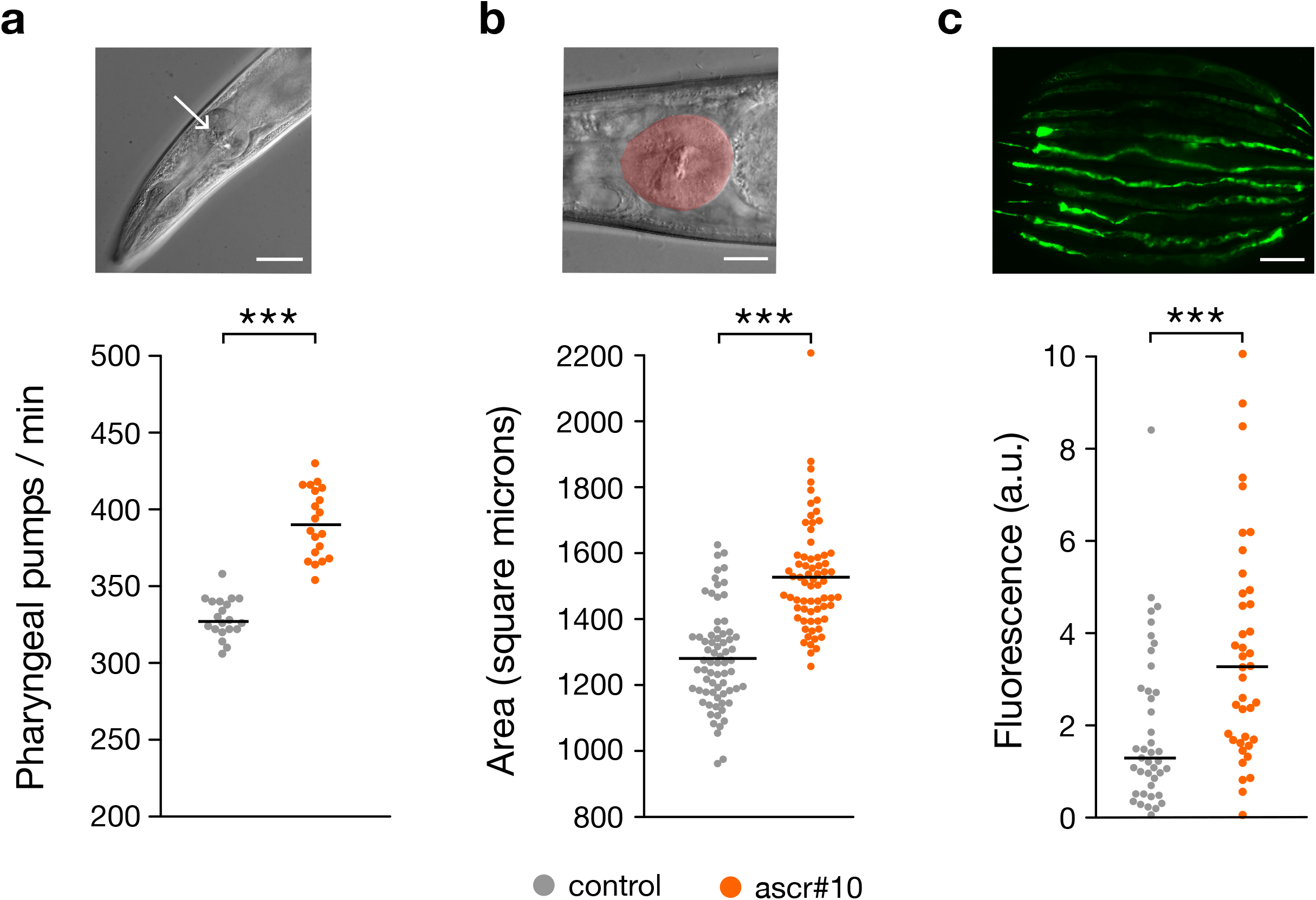
The effects of ascr#10 on pharyngeal function and morphology. **a**, DIC image of the pharynx and rate of pharyngeal pumping in young adult hermaphrodites exposed to control or ascr#10 for one hour (n = 20). The arrow points to the grinder within the posterior bulb of the pharynx. Scale bar = 40 *μ*m. **b**, Area of the posterior bulb (highlighted by false coloring) in hermaphrodites (Day 8 of adulthood) raised on control or ascr#10 (n = 70 for control and n = 68 for ascr#10). Scale bar = 20 *μ*m. **c**, Representative image and quantification of fluorescence in hermaphrodites (Day 8 of adulthood) raised on control or ascr#10 and fed OP50 *E. coli* expressing GFP (n = 41 for control and n = 40 for ascr#10). Scale bar = 200 *μ*m. In all three panels, p-values are from Kolmogorov-Smirnov test; ***, P < 0.01.

### Broad effects of ascr#10 on reproduction-related traits and longevity

There are five notable aspects of the response of *C. elegans* hermaphrodites to brief exposure to physiologically relevant (Aprison and Ruvinsky 2017) concentrations of the male pheromone ascr#10.

First, several thousand genes constituting over 1/4 of the detected genes showed significant change in expression, although nearly all by less than two-fold. Therefore, the physiological and behavioral effects of ascr#10 stem from modest fine-tuning of multiple genetic programs, not from substantial changes in a few select pathways. Broad changes in gene expression upon encountering the complete male pheromone bouquet were recently reported (Booth, et al. 2022).

Second, the most conspicuous transcriptomic changes induced by ascr#10 were the upregulation of the oogenic gene expression and the downregulation of the spermatogenic gene expression, demonstrating that a single component of the male pheromone is sufficient to feminize germline gene expression programs. Simultaneous hermaphrodites, such as *C. elegans*, must balance demands of sperm and oocyte production in the same organism (Charnov 1979). The male pheromone ascr#10 signals to hermaphrodites the availability of male sperm and thus reduced need for self-sperm. We interpret the ensuing transcriptomic changes as tuning gene expression programs to optimize oogenesis, in line with the hypothesis in Figure 1a. We are not aware of an experimental demonstration of such a phenomenon in any species. Theory predicts that selection should favor deployment of reproductive resources by hermaphrodites (between male and female functions) that best reflects mating opportunities (Charnov 1982). Experimental evidence supports this idea. For example, hermaphrodites of the plant *Mercurialis annua* mating in the absence of males evolved greater male allocation (Dorken and Pannell 2009). Our results are consistent with feminization of the hermaphrodite germline transcriptome in the presence of a male pheromone.

Third, in our experiments, young adult hermaphrodites were exposed to the pheromone several hours after the switch between spermatogenesis and oogenesis that occurs during the last larval stage (Kimble and Crittenden 2007). It may seem surprising that animals that apparently irreversibly committed to oocyte production continue to express many genes associated with sperm production. Yet, transcriptome profiling of young adult hermaphrodites similar in age to the ones we used in this study shows substantial continued expression of genes associated with spermatogenesis (Ortiz, et al. 2014; Ebbing, et al. 2018). These results indicate that the binary switch in the type of produced gametes is undergirded by a more gradual change in gene expression. A switch from spermatogenesis to oogenesis is thought to involve release of post-transcriptional repression of oogenic transcripts (Noble, et al. 2016). Understanding how (or whether) this process is related to the male pheromone-induced bias of the pool of mRNAs toward oogenic profile may be instructive for understanding germline development in *C. elegans*.

Fourth, *C. elegans* hermaphrodites continue to respond to male pheromones even though males and mating are relatively rare in natural populations (Frezal and Felix 2015). Why is this relic of the ancestral gonochoristic mode of reproduction (Cutter, et al. 2019) retained in a hermaphroditic species? In natural habitats worms commonly experience stress conditions that increase the frequency of males. The resulting outcrossing is favored and advantageous under such conditions (Morran, Cappy, et al. 2009; Morran, Parmenter, et al. 2009). We propose that increased reproductive success conferred by ascr#10 in challenging environments contributes to the retention of this male pheromone in a hermaphroditic species. Consistent with this idea, ascr#10 facilitates reproductive recovery from heat stress (Aprison and Ruvinsky 2015).

Finally, interactions with males are detrimental to *C. elegans* hermaphrodites (Shi and Murphy 2014; Ting, et al. 2014; Booth, et al. 2022) and some of this effect is due to male-excreted compounds (Maures, et al. 2014; Shi, et al. 2017). Previously, we showed that ascr#10 shortens the hermaphrodite lifespan (Aprison and Ruvinsky 2016; Ludewig, et al. 2019). The physiological changes induced by male pheromones that reduce longevity are only beginning to be understood. Our work has identified three plausibly detrimental effects of ascr#10 that likely contribute to the shortening of the hermaphrodite lifespan. 1) Pharyngeal hypertrophy and the associated persistent bacterial infections (Figure 2). 2) Increased yolk production in hermaphrodites (Figure 1d) can result in senescent pathologies (Garigan, et al. 2002; Zimmerman, et al. 2015; Seah, et al. 2016; Ezcurra, et al. 2018) and, in general, yolk production has opposite effects on reproductive success and longevity (Wu, et al. 2022). 3) Physiological cell death in the germline is increased by ascr#10 and is required for improving oocyte quality (Aprison, et al. 2022), but it may result in gonad pathologies (de la Guardia, et al. 2016). In essence, the male pheromone promotes investment in the oogenic germline in youthful hermaphrodites at the cost of accelerated post-reproductive senescence.

## Materials and methods

### Analysis of differential gene expression in presence of ascr#10

We used the data from (Aprison, et al. 2022) to carry out all analyses (GSE193636). Briefly, wild type N2 hermaphrodites were synchronized and exposed to ascr#10 ∼3-4 hours after the L4-to-adult molt (48 hours post release from L1 arrest). Germline production switches from spermatogenesis to oogenesis during the L4 stage (Kimble and Crittenden 2007). Worms were harvested for RNA preparation either ∼3-4 hours before the onset of reproduction (50 hours post release from L1 arrest) or ∼3-4 hours after the onset of egg laying (58 hours post release from L1 arrest).

We restricted analyses only to those genes that were identified across both conditions. Consequently, the number of differentially expressed genes in our analyses is smaller than previously reported (1,667 and 3,627 reported originally in pre-reproductive and reproductive animals in response to ascr#10 vs 1,663 and 3,622 reported here), but the fold-changes and q-values are identical with Aprison *et al*. (Aprison, et al. 2022).

All code required to completely reproduce our analyses is available in GitHub [https://github.com/dangeles/Ascr10xomics]. Statistical functions and large pieces of code were written as python v.3.7(van Rossum 2009) scripts that can be loaded locally; the analyses are carried out in annotated Jupyter notebooks (Granger and Pérez 2021) which were used to generate the supplementary information (Supplementary Information).

We tested tissues for directional enrichment using a binomial test. We downloaded the complete tissue expression annotations from WormBase (Davis, et al. 2022). We restricted our analysis to tissues that had at least 5 annotated genes. We did not use genes if they had promiscuous tissue expression–we dropped any genes that were annotated in 30 or more terms. If a term had annotated genes that were all considered promiscuous, that term was also dropped. We tested terms for regulatory enrichment by using a binomial test, where the proportion of genes expected to increase expression was given by the net fraction of genes that were upregulated in each condition, followed by a Benjamini-Hochberg correction. Exclusively for visualization purposes, we removed redundant tissue terms by identifying terms that were more than 75% similar by Jaccard similarity and removing the term with fewer annotated gene expression levels. Tissues were concordant between the two compared timepoints (50 and 58 hours post release from the L1 arrest), though q-values were lower post-onset of reproduction.

For our analyses of the single-cell gene expression atlas (Taylor, et al. 2021) using SCANPY (Wolf, et al. 2018), we removed all cells annotated ‘Unknown’ and ‘Unannotated’. We also removed ‘Muscle_mesoderm’ cells because they clustered apart from the remaining cell types and distorted the UMAP diagrams; however, removing these cells did not alter our conclusions. We filtered cells that showed expression of fewer than 150 genes and removed any genes expressed in fewer than 50 cells. We removed cells that had fewer than 250 total counts, mitochondrial content <1% or >10%, riboprotein content <1% or >20%, and NADH enzyme content <.3% or >3%. If a cell was annotated as a neuron, we removed cells that had < 800 total counts, because we observed that neurons had much greater read content than the other cell types. Given the low read content, we normalized counts in cells to counts per thousand, as suggested previously (Lun 2018).

To generate UMAPs, we subsampled the dataset to equally represent all tissues with 2000 cells (or less if fewer cells were annotated in the dataset). This allowed the UMAP to show better separation among tissues. We reduced the dimensionality of the dataset using PCA and keeping the first 15 principal components. Next, we computed the 100 nearest neighbors per cell, and computed the UMAP using SCANPY’s UMAP function, initializing the UMAP with pre-computed PAGA positions (Wolf, et al. 2019). Next, we computed signature enrichment scores. To generate an ascr#10-upregulated (or downregulated) signature, we identified the genes that were differentially expressed at 50 and 58 hours (on vs. off ascr#10), that changed expression in the same direction, and that increased (decreased) expression in response to ascr#10. For comparison, we also generated a signature with 159 randomly selected genes. We calculated an enrichment score using SCANPY’s ‘score_genes’ function. This function reproduces the score derived in (Satija, et al. 2015), which works by computing the average expression level of the input signature genes and subtracting a random reference set of genes. The random reference set is chosen so that the distribution of average gene expression levels between the input and reference sets are similar.

### Measurements of pharyngeal phenotypes in hermaphrodites exposed to ascr#10

Concentrated ascr#10 was stored in ethanol at −20^°^C. This stock was diluted with water and a total of 100μL of ascaroside solution containing 2.2 femtograms per plate was applied to the plate and spread evenly with a sterile bent glass rod. Plates were incubated at 20^°^C overnight to allow the ascaroside to be absorbed. Control plates were prepared in the same manner using water as the control. The following day the plates were seeded with 20μL of a 1:10 dilution of OP50 overnight culture and allowed to grow overnight at 20^°^C before use. All experiments were conducted using wild type N2 hermaphrodites.

To measure pharyngeal pumping rates, hermaphrodites were synchronized by hypochlorite treatment and tested at ∼70 hours post release from L1 arrest (Day 2 adults). Ten hermaphrodites per plate were transferred to either control plates or ascr#10 plates prepared as above and allowed to acclimate for one hour. The number of pharyngeal pumps was counted for 20 seconds. This was done 3X for each worm waiting at least 20 seconds between counts.

To measure the area of the terminal bulb of the pharynx in worms that had been raised on control plates or ascr#10 plates, Day 8 adults were mounted on 2% agarose pads and images were recorded using a Leica DM5000B microscope fitted with a Retiga 2000R camera and quantified using ImageJ.

To measure intestinal colonization by bacteria, hermaphrodites were grown on either control or ascr#10 plates seeded with OP50 *E. coli* carrying the pFPV25.1 plasmid that encodes GFP. Worms were transferred every other day until they were Day 8 adults. Day 8 adults were mounted on 2% agarose pads and the slides were imaged on a Leica DM5000B microscope using a Retiga 2000R camera. Images were stitched together using the ImageJ plug-in (Preibisch, et al. 2009). The entire intestine was outlined in each hermaphrodite and the corrected total fluorescence for each organ was determined in ImageJ. Since Day 8 adults exhibit increased autofluorescence (Teuscher and Ewald 2018), Day 8 adults raised on control or ascr#10 plates seeded with regular OP50 were also compared and the amount of autofluorescence was quantified as above.

## Acknowledgements

We thank R. Morimoto for generous hospitality and F. Schroeder for ascr#10. This work was funded in part by the NIH (R01GM126125) grant to I.R. We thank WormBase and the Caenorhabditis Genetics Center (CGC). WormBase is supported by grant U41 HG002223 from the National Human Genome Research Institute at the NIH, the UK Medical Research Council, and the UK Biotechnology and Biological Sciences Research Council. The CGC is funded by the NIH Office of Research Infrastructure Programs (P40 OD010440).

