## Supplemental Information for "A *C. elegans* male pheromone feminizes germline gene expression in hermaphrodites and imposes life-history costs"

### 01_BasicDataExploration

September 27, 2022

#### 1 Data exploration and data wrangling

For all the notebooks you will find here, we first load the same set of libraries, and we also load the tidy data into a variable called `res`.

[1]:
```
import pandas as pd
import numpy as np
import scipy
import matplotlib as mpl
import matplotlib.pyplot as plt
import seaborn as sns
from matplotlib import rc

rc('text', usetex=False)
rc('text.latex', preamble=r'\usepackage{cmbright}')
rc('font', **{'family': 'sans-serif', 'sans-serif': ['Helvetica']})

%matplotlib inline

# This enables SVG graphics inline.
%config InlineBackend.figure_formats = {'png', 'retina'}

rc = {'lines.linewidth': 2,
      'axes.labelsize': 25,
      'axes.titlesize': 25,
      'axes.facecolor': 'DFDFE5'}
sns.set_context('notebook', rc=rc)
sns.set_style("dark")

mpl.rcParams['xtick.labelsize'] = 18
mpl.rcParams['ytick.labelsize'] = 18
mpl.rcParams['legend.fontsize'] = 20

res = pd.read_csv('../data/master_table.tsv', sep='\t')
print('Number of genes included in analysis:', len(res))
print('DE genes that will be analyzed at 50 hrs: ', (res['padj-50'] < 0.05).
 ↪sum())
```

```
print('DE genes that will be analyzed at 58 hrs: ', (res['padj-58'] < 0.05).
 ↪sum())
print('DE genes in both conditions: ', res.query('`padj-50` < 0.05 & `padj-58`␣
 ↪< 0.05').shape[0])
```

```
Number of genes included in analysis: 12244
DE genes that will be analyzed at 50 hrs:  1663
DE genes that will be analyzed at 58 hrs:  3622
DE genes in both conditions:  741
```

#### 2 Data Exploration and QC

Briefly, we performed RNA-seq of animals exposed to ascr#10 until they were pre-licensed adults or until they were licensed adults (recall the licensing event is egg-laying); this corresponds to animals aged 50hrs and 58hrs respectively. In a previously published article, we processed this data using Salmon + DESeq to identify differentially expressed genes.

[2]:
```
res = res[(res['padj-50'] < 0.05) | (res['padj-58'] < 0.05)]

def save(name, transparent=False):
    """Wrapper to abbreviate save file location."""
    plt.savefig('../figs/' + name, bbox_inches='tight', transparent=transparent)

sns.relplot(
    data=res,
    x="log2FoldChange-50", y="log2FoldChange-58",
    size='neglogq-58',
    kind="scatter", sizes=(25, 300), hue='Sign-WT',
    palette=['black', 'tab:red'], hue_order=['Same', 'Different'],
    alpha=0.5
)

print('Number of genes that have the same log(FC) sign and are DE in at least␣
 ↪one condition:')
print(res.query('`Sign-WT` == "Same" & (`padj-58` < 0.05 | `padj-50` < 0.05)' ).
 ↪shape[0])
print('Number of genes that have different log(FC) sign and are DE in at least␣
 ↪one condition:')
print(res.query('`Sign-WT` != "Same" & (`padj-58` < 0.05 | `padj-50` < 0.05)' ).
 ↪shape[0])
```

```
Number of genes that have the same log(FC) sign and are DE in at least one
condition:
3087
Number of genes that have different log(FC) sign and are DE in at least one
condition:
1457
```

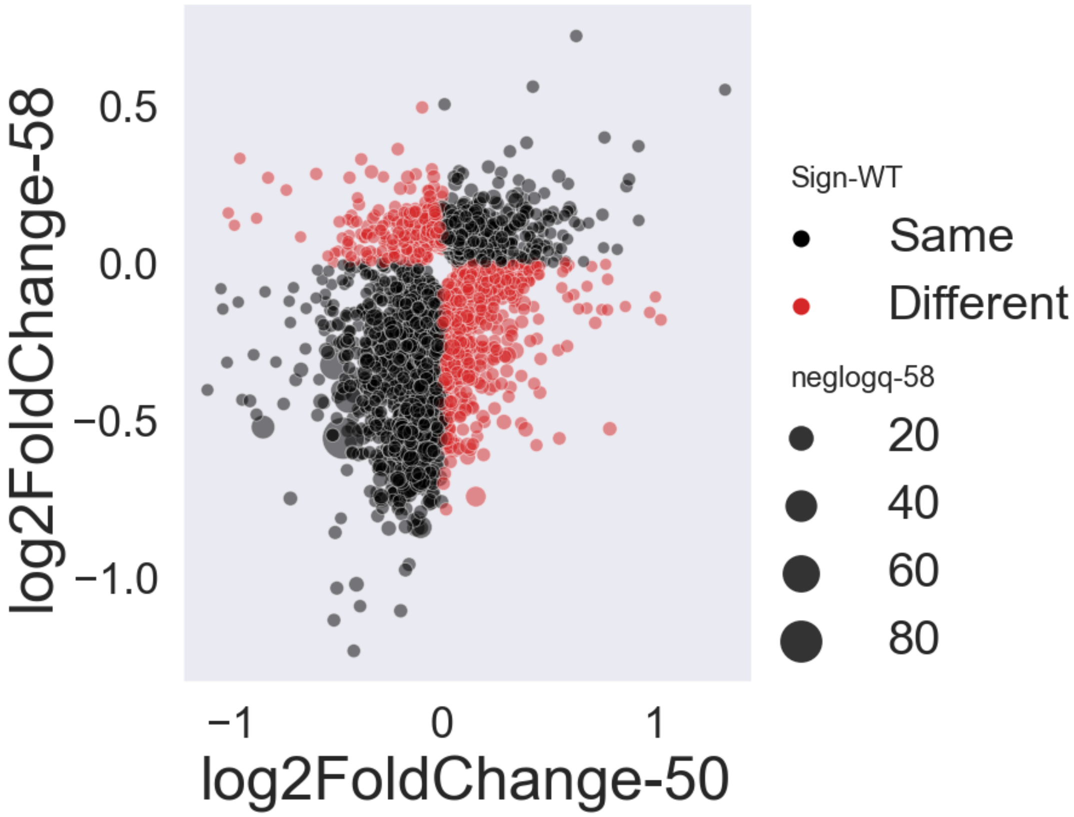


Perhaps a more productive view of the data comes from looking at the genes that are commony differentially expressed in response to ascr#10 exposure at both timepoints:

[3]:
```
sns.relplot(
    data=res[(res['padj-58'] < 0.05) & (res['padj-50'] < 0.05)],
    x="log2FoldChange-50", y="log2FoldChange-58",
    size='neglogq-58',
    kind="scatter", sizes=(25, 300), hue='Sign-WT',
    palette=['black', 'tab:red'], hue_order=['Same', 'Different'],
    alpha=0.75,
)

plt.plot(res['log2FoldChange-50'], 1 * res['log2FoldChange-50'],
         label='50hrs = 58hrs', color='blue', ls='--')

print('Number of genes that have the same log(FC) sign and are DE in at least␣
 ↪one condition:')
print(res.query('`Sign-WT` == "Same" & (`padj-58` < 0.05 & `padj-50` < 0.05)' ).
 ↪shape[0])
print('Number of genes that have different log(FC) sign and are DE in at least␣
 ↪one condition:')
```

```
print(res.query('`Sign-WT` != "Same" & (`padj-58` < 0.05 & `padj-50` < 0.05)' ).
↪shape[0])
```

```
Number of genes that have the same log(FC) sign and are DE in at least one
condition:
650
Number of genes that have different log(FC) sign and are DE in at least one
condition:
91
```

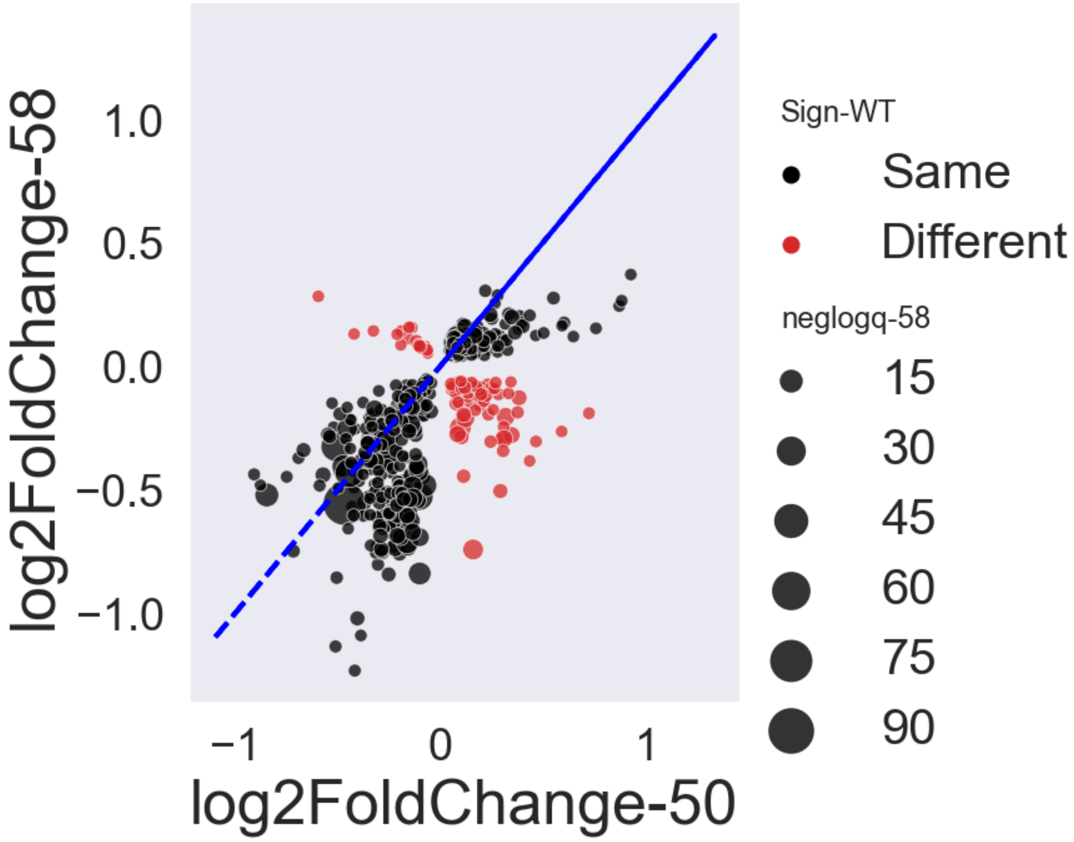


In the above graph, it becomes clear that, at least for commonly regulated genes, the majority of genes are positively correlated between both conditions. The blue line in the graph is the line $y = x$, suggesting that, on average, ascr#10 responsive genes do not change their response in a time-dependent manner.

Moreover, not that the off-diagonal points are much, much fewer than before! This suggests that those off-diagonal genes are false measurements in at least one condition. A slightly more rigorous argument follows:

Consider those genes that are DE in at least one condition (the **Union**), and which are either correlated or anticorrelated. Assume that the incomplete overlap between both conditions is due to false positives and false negatives, and is NOT the result of additional layers of regulation (a major, but reasonable assumption). Then, it follows that genes DE in BOTH conditions (the **Intersection**) are genes whose measurements were more replicable on average, since they were detected in both conditions.

Then, it follows that the proportion of correlated : anti-correlated genes should be the same if we measure it in *either* the union or the intersection. In the **union**, this proportion is $1455/3084 \sim .5$, whereas in the **intersection**, the proportion is $91/650 = .14$. The fraction of anti-correlated genes has gone down significantly. *Therefore, under the assumption that the genetic regulation is identical at 50 AND at 58 hours, we conclude that the off-diagonal genes are anticorrelated due to noise, and we ignore any genes in the* ***intersection*** *that are anti-correlated.*

### 02_TissueEnrichment

September 27, 2022

#### 1 Tissue-specificity of the ascr#10 response

In this notebook, we explore the tissue specificity of the ascr#10 response in *C. elegans*. Please note that most enrichment analyses first count the number of DE genes associated with a specific term, and compare that number with the expected number under a uniform random model–this is called a hypergeometric test. In this notebook, we do NOT perform hypergeometric tests to identify enriched tissues. We take a different approach.

In this notebook, we download the gene expression patterns from Wormbase, and then we test whether genes that are expressed in a given tissue are going UP(down) more often than expected by random chance. We do this using a binomial test.

The WormBase gene expression-tissue database has been processed in a fairly specific way. First, we removed all non-DE genes at 50 or at 58hrs from the database. Next, we dropped all tissues that did not have at least 5 genes annotated to them. Finally, we removed 'promiscuous genes', genes that are annotated to many (>30) tissues. Originally, the database had 1,535 tissues and after processing we have 264 tissues.

[1]:
```
import sys
sys.path.append('../python')
import tissue_utils as utils  # the functions I wrote are here
import json
import pandas as pd
import numpy as np
import scipy
import matplotlib as mpl
import matplotlib.pyplot as plt
import seaborn as sns

from statsmodels.stats.multitest import fdrcorrection

from matplotlib import rc
rc('text', usetex=False)
# rc('text.latex', preamble=r'\usepackage{cmbright}')
rc('font', **{'family': 'sans-serif', 'sans-serif': ['Helvetica']})

%matplotlib inline

# This enables SVG graphics inline.
```

```
%config InlineBackend.figure_formats = {'png', 'retina'}

rc = {'lines.linewidth': 2,
      'axes.labelsize': 25,
      'axes.titlesize': 25,
      'axes.facecolor': 'DFDFE5'}
sns.set_context('notebook', rc=rc)
sns.set_style("dark")

mpl.rcParams['xtick.labelsize'] = 18
mpl.rcParams['ytick.labelsize'] = 18
mpl.rcParams['legend.fontsize'] = 20
# load stuff:
res = pd.read_csv('../data/master_table.tsv', sep='\t', index_col=0)

cols = ['externalgenename', 'log2FoldChange-{0}', 'padj-{0}']
fetch = lambda x: [c.format(x) for c in cols]
rename = lambda x: {'log2FoldChange-{0}'.format(x): 'log2FoldChange',␣
 ↪'padj-{0}'.format(x): 'padj'}

# data wrangling to create a new dataframe that contains the `results` data,␣
 ↪but will also include
# the tissue annotations
tmp_list = []
for suffix in ['50', '58']:
    tmp = res[fetch(suffix)].rename(columns=rename(suffix))
    tmp['data'] = suffix
    tmp = tmp[tmp.padj < 0.05]
    print(suffix, tmp.shape)
    tmp_list += [tmp]

annotated_data = pd.concat(tmp_list)
cat_type = pd.CategoricalDtype(categories=['50', '58'], ordered=True)
annotated_data.data = annotated_data.data.astype(cat_type)

# load tissue dictionary:
tissues = utils.load_tissues(res.index, 5, 30)
```

```
50 (1663, 4)
58 (3622, 4)
tissues originally: 2359
tissues afterwards: 846
```

[2]:
```
counts = annotated_data[annotated_data.padj < 0.05].groupby('data').
 ↪externalgenename.count().rename('DEG count').reset_index()
sns.stripplot(y='data', x='DEG count', hue='data', data=counts,
```

```
              s=15, palette={'50': 'tab:blue', '58': 'tab:green', 'tph1-50':␣
↪'blue', 'tph1-58': 'green', 'pqm1': 'tab:purple'})
plt.legend([])
plt.xlabel('Number of DEGs')
plt.savefig('../figs/DEG_number.svg', bbox_inches='tight')
```

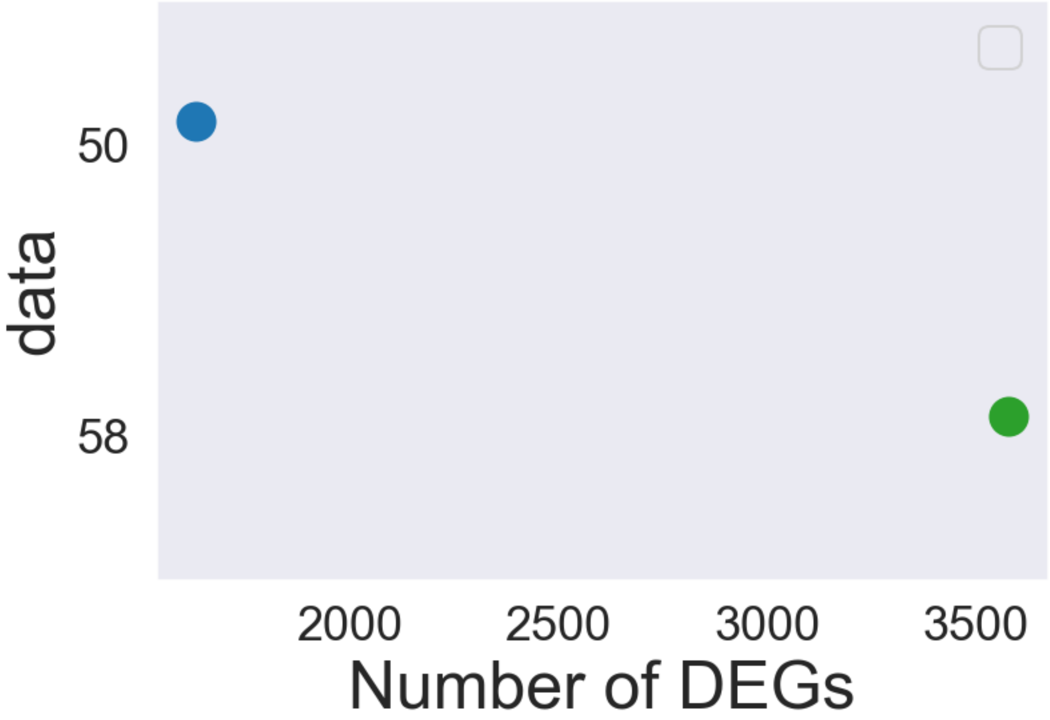


##### 1.1 Testing the tissues.

In the cell above, we loaded our data, and we also loaded our gene expression pattern database into a variable called `tissues`. Here, we perform a binomial test on each of the tissue programs.

[3]:
```
# perform a binomial test for change in direction:
alpha = 0.05
data = utils.test_tissue_direction(res[(res['padj-58'] < 0.05) &␣
↪(res['Sign-WT'] == 'Same')], tissues)
data_50 = utils.test_tissue_direction(res[(res['padj-50'] < 0.05) &␣
↪(res['Sign-WT'] == 'Same')], tissues, col='log2FoldChange-50')

data = pd.concat([data, data_50], keys=['58hrs', '50hrs']
                ).reset_index().rename(
                columns={'level_0': 'data'}
                ).drop('level_1', axis=1)
```

```
data = utils.fdr_correct(data)
data.sort_values('fdr', inplace=True)

data['MeanFracPos'] = data.groupby(['tissue', 'data']).FracPos.transform(np.
 ↪mean)
data['MeanFracPos'] = data.groupby(['tissue', 'data']).FracPos.transform(np.
 ↪mean)

# keep only significant results:
keep = data.groupby('tissue').sig.sum()
data = data[data.tissue.isin(keep[keep > 0].index)]

m = 'There were {0} tissues that changed more in one direction than expected by␣
 ↪random chance in dataset {1}'
for n, g in data.groupby('data'):
    print(m.format(len(g), n))

data.head()
```

```
There were 41 tissues that changed more in one direction than expected by random
chance in dataset 50hrs
There were 44 tissues that changed more in one direction than expected by random
chance in dataset 58hrs
```

[3]:

```
    data                  tissue          pval   FracPos  FracPosExpected  \
0  58hrs               germ line  2.896386e-23  0.951613         0.646437
1  58hrs                    Cell  4.057242e-11  0.936842         0.646437
2  58hrs          nervous system  6.766480e-11  0.836066         0.646437
3  58hrs  body wall musculature  1.081277e-10  0.817568         0.646437
4  58hrs                 pharynx  9.193526e-10  0.797688         0.646437

            fdr    neglogq   sig  MeanFracPos
0  7.125109e-21  20.147208  True     0.951613
1  4.990408e-09   8.301864  True     0.936842
2  5.548514e-09   8.255823  True     0.836066
3  6.649853e-09   8.177188  True     0.817568
4  4.523215e-08   7.344553  True     0.797688
```

Next, let's remove uninformative terms like `cell`, `tail` or `head`. I will also remove highly similar terms to prevent too much redundancy. For completeness, I've made sure to print all tissues that I will remove:

[4]:

```
remove = ['Cell', 'tail', 'head', 'male gonad', 'gonad', ]  # so broad. why␣
 ↪ever have these terms...
remove = utils.similarity_trimming(data.tissue.unique(), tissues, remove)
print('The following list of tissues will be removed from the plot',␣
 ↪list(set(remove)))
```

```
The following list of tissues will be removed from the plot ['mu-int-R', 'vulval
```

```
muscle', 'male gonad', 'hermaphrodite gonad', 'tail', 'nervous system', 'ventral
cord neuron', 'P0', 'tail neuron', 'mu-int-L', 'anal depressor muscle', 'body
region', 'Z2', 'gonad', 'anterior distal tip cell', 'Cell', 'head', 'Psub2']
```

Let's plot the results:

[5]:
```
to_plot = data[(~data.tissue.isin(remove)) & (data.sig == True)].head(15)  #␣
↪get top 15 hits or so
to_plot = data[(data.tissue.isin(to_plot.tissue))]  # only use the top 10-15␣
↪tissues for this plot

tissues_plotted = to_plot.tissue.unique()
fig, ax = utils.pretty_GSEA_plots(to_plot, sig_level=alpha, size=20, alpha=0.6)
ax.yaxis.grid(color='black', linewidth=1)
ax.set_xlabel('-log(q)', fontsize=30)
plt.savefig('../figs/tissue_GSEA.svg', bbox_inches='tight', transparent=False)
```

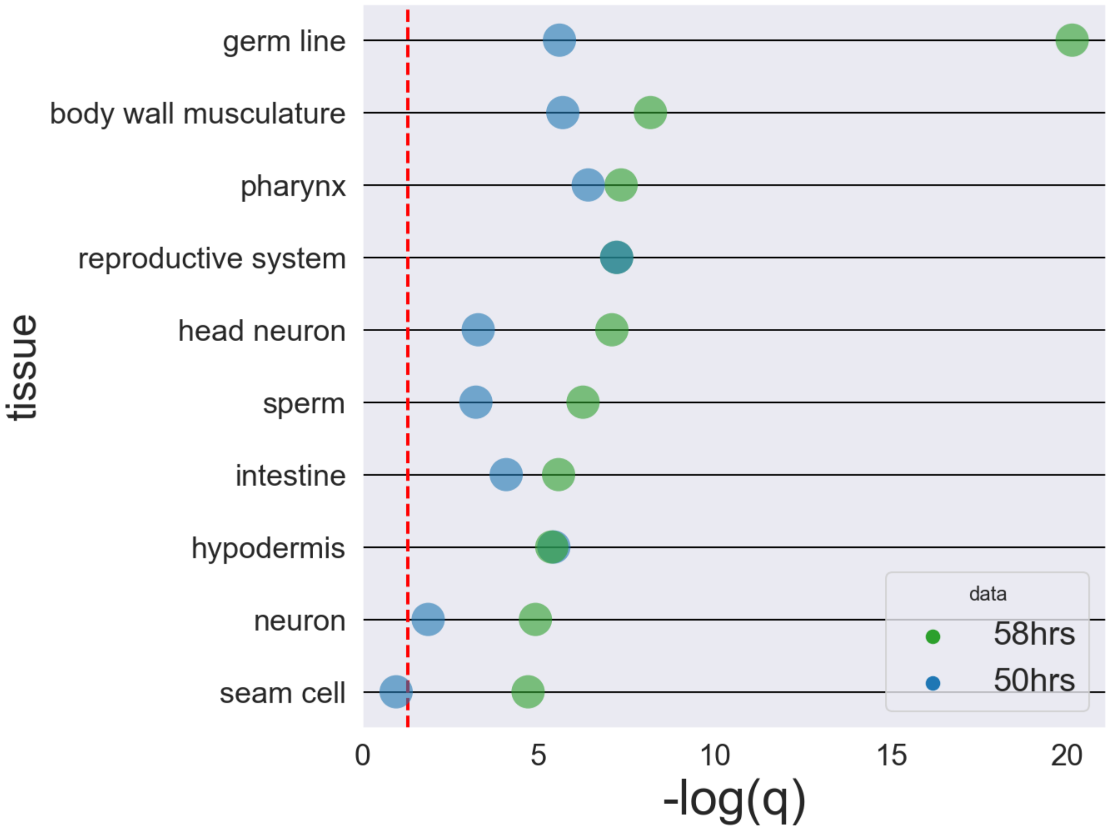


#### 1.2 Volcano Plot

Here is the volcano plot that was reproduced in the paper:

```
[6]: to_plot['Excess'] = (to_plot.FracPos - to_plot.FracPosExpected) / to_plot.
     ↪FracPosExpected

     fig, ax = plt.subplots(figsize=(8, 8))
     sns.scatterplot(x='Excess', y='neglogq', data=to_plot[to_plot.data  ==␣
     ↪'58hrs'], s=150, color='tab:green', ax=ax)

     for tissue, df in to_plot[to_plot.data == '58hrs'].groupby('tissue'):
         plt.annotate(tissue, (df.Excess.unique()[0], df.neglogq.unique()[0]),
                      fontsize=20)
     #     print(df.head())

     # plt.axvline(to_plot.FracPosExpected.unique()[0], color='red', ls='--',␣
     ↪label='Expected % of Genes UP')
     plt.axhline(1, color='black', ls='-.', label='Expected % of Genes UP')
     plt.axvline(0, color='red', ls='--')
     plt.xlim(-1, 1)
     plt.xlabel('Excess Positivity over Expectation')
     plt.ylabel('-log$_{10}$(q)')

     plt.savefig('../figs/tissue_volcano.svg', bbox_inches='tight',␣
     ↪transparent=False)
```

```
/Users/davidangeles/opt/anaconda3/lib/python3.7/site-
packages/ipykernel_launcher.py:1: SettingWithCopyWarning:
A value is trying to be set on a copy of a slice from a DataFrame.
Try using .loc[row_indexer,col_indexer] = value instead

See the caveats in the documentation: https://pandas.pydata.org/pandas-
docs/stable/user_guide/indexing.html#returning-a-view-versus-a-copy
  """Entry point for launching an IPython kernel.
```

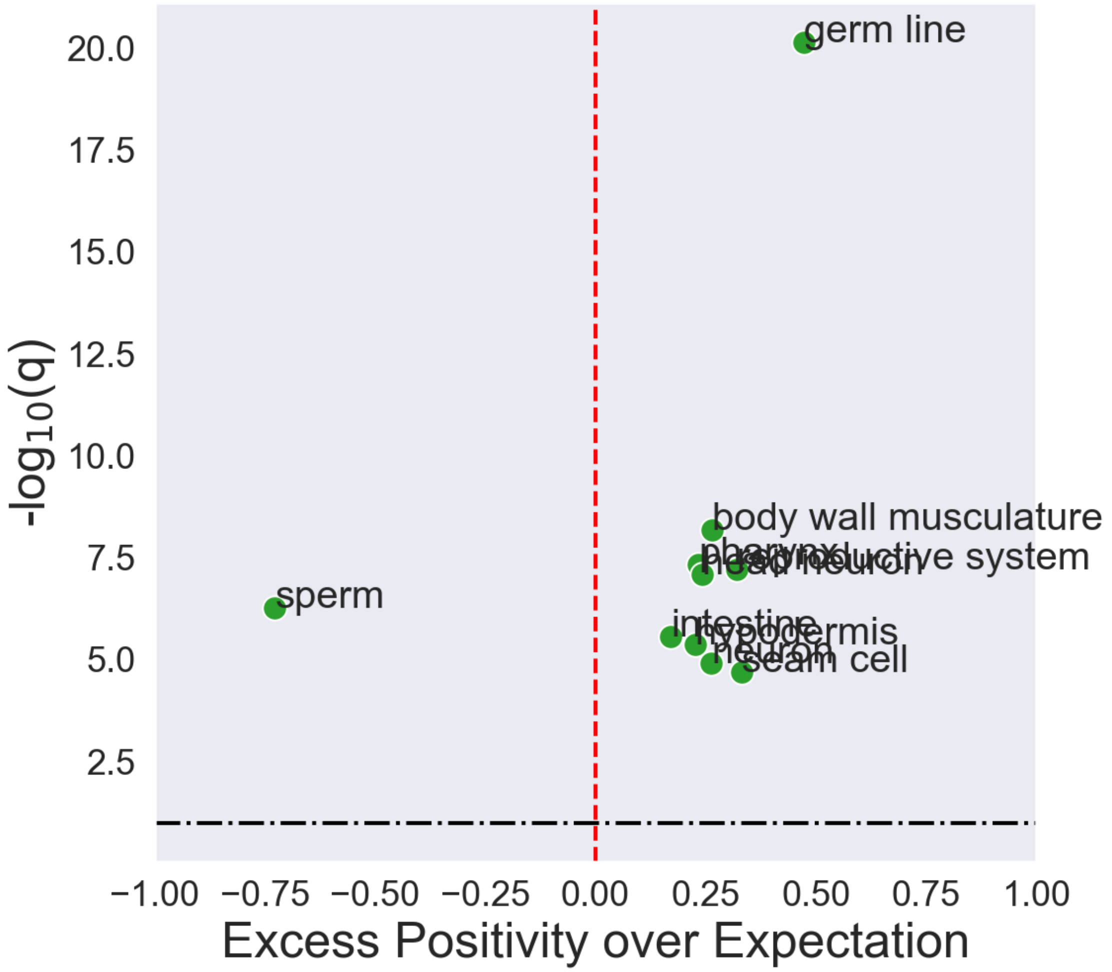


#### 2 Directional analysis of gene subsets.

In the next section, we pull out all the vitellogenin genes, the major sperm protein genes and all ribosome small / large subunit genes, and we plot their measured log Fold Change at 50 and 58 hours.

[7]:
```
res = pd.read_csv('../data/master_table.tsv', sep='\t', index_col=0)
res = res.dropna(subset=['log2FoldChange-50', 'log2FoldChange-58'])
tissues = utils.load_tissues(res.index, 5, 30)
annotated_data = res.join(tissues.set_index('wbid')).dropna(subset=['tissue'])
```

```
tissues originally: 2359
tissues afterwards: 846
```

[8]:
```
def plot_gene_fam(res, title, ax=ax):
    """Plot genes that start with the suffix `title` in a stripplot"""
```

```
    selected = (res.Gene.fillna('').str.contains(title)) &\
               (res.Gene.str.split('-', expand=True)[0].astype(str).
↪apply(lambda x: len(x)) == 3)

    colors = {'58': 'tab:green', '50': 'tab:blue', 'pqm1': 'tab:purple',
              'tph1-50': 'blue', 'tph1-58': 'green'}

    log_cols = ['Gene'] + [c for c in res.columns if ('log2' in c)]
    pval_cols = ['Gene'] + [c for c in res.columns if ('padj' in c)]

    tidy = res[selected][log_cols].melt(id_vars='Gene', var_name='Condition',␣
↪value_name='logFoldChange')
    tidy_qvals = res[selected][pval_cols].melt(id_vars='Gene',␣
↪var_name='Condition', value_name='qval')

    tidy.Condition = tidy.Condition.str.replace('log2FoldChange-', '')
    tidy_qvals.Condition = tidy_qvals.Condition.str.replace('padj-', '')

    tidy = tidy.join(tidy_qvals.set_index(['Gene', 'Condition']), on=['Gene',␣
↪'Condition'])
    tidy['Sig'] = tidy.qval < 0.1

    print(tidy.Gene.unique())
    if len(tidy) > 30:
        alpha = .3
    else:
        alpha= 1

    sns.stripplot(y='Condition', x='logFoldChange', hue='Condition',␣
↪alpha=alpha,
                  s=8, data=tidy, orient='h', palette=colors, ax=ax)
    ax.axvline(0, color='black', ls='--', lw=1)
    ax.legend(loc=(1, .3), title='Diff Expressed')
    ax.set_title(title + ' genes')

def three_panel_plot(res):
    """A function to plot the three desired panels"""
    fig, ax = plt.subplots(ncols=3, sharey=True, sharex=False, figsize=(15, 2.
↪5))
    plot_gene_fam(res, title='vit', ax=ax[0])
    plot_gene_fam(res, title='msp', ax=ax[1])
    plot_gene_fam(res, title='^rp[sl]', ax=ax[2])

    ax[2].set_title('rps/rpl genes')

    ax[0].legend([])
```

```
    ax[1].legend([])

    ax[0].set_xlabel('')
    ax[2].set_xlabel('')

    ax[1].set_xlabel('log$_2$(Fold Change)')

    ax[0].set_xlim(-.7, .7)
    ax[1].set_xlim(-.7, .7)
    ax[2].set_xlim(-.2, .2)
    return fig, ax


fig, ax = three_panel_plot(res)
plt.savefig('../figs/gene_programs.svg', bbox_inches='tight', transparent=False)
```

```
['vit-1' 'vit-2' 'vit-3' 'vit-4' 'vit-5' 'vit-6']
['msp-3' 'msp-19' 'msp-31' 'msp-33' 'msp-36' 'msp-38' 'msp-40' 'msp-45'
 'msp-49' 'msp-50' 'msp-51' 'msp-53' 'msp-55' 'msp-56' 'msp-57' 'msp-59'
 'msp-64' 'msp-65' 'msp-76' 'msp-77' 'msp-78' 'msp-81' 'msp-113' 'msp-142'
 'msp-152']
['rpl-1' 'rpl-2' 'rpl-3' 'rpl-4' 'rpl-5' 'rpl-6' 'rpl-7' 'rpl-7A' 'rpl-9'
 'rpl-10' 'rpl-11.1' 'rpl-11.2' 'rpl-12' 'rpl-13' 'rpl-14' 'rpl-15'
 'rpl-16' 'rpl-17' 'rpl-18' 'rpl-19' 'rpl-20' 'rpl-21' 'rpl-22' 'rpl-23'
 'rpl-24.1' 'rpl-24.2' 'rpl-25.1' 'rpl-25.2' 'rpl-26' 'rpl-27' 'rpl-28'
 'rpl-29' 'rpl-30' 'rpl-31' 'rpl-32' 'rpl-33' 'rpl-34' 'rpl-35' 'rpl-36'
 'rpl-37' 'rpl-38' 'rpl-39' 'rpl-41' 'rpl-43' 'rps-0' 'rps-1' 'rps-2'
 'rps-3' 'rps-4' 'rps-5' 'rps-6' 'rps-7' 'rps-8' 'rps-9' 'rps-10' 'rps-11'
 'rps-12' 'rps-13' 'rps-14' 'rps-15' 'rps-16' 'rps-17' 'rps-18' 'rps-19'
 'rps-20' 'rps-21' 'rps-22' 'rps-23' 'rps-24' 'rps-25' 'rps-26' 'rps-27'
 'rps-28' 'rps-29' 'rps-30']
```

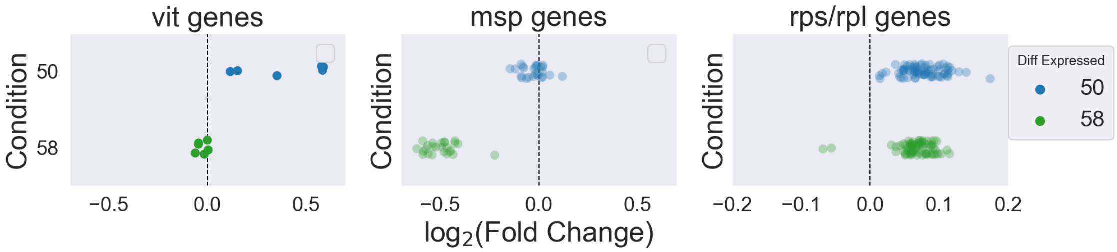

### 03_Singlecell-Analysis

February 16, 2023

#### 1 Re-analysis of the Taylor *et al.* single-cell dataset for enrichment of ascr#10 signatures.

[1]:
```
import pandas as pd
import numpy as np
import matplotlib as mpl
import matplotlib.pyplot as plt
import seaborn as sns
from matplotlib import rc
import scanpy as sc

rc('text', usetex=False)
rc('text.latex', preamble=r'\usepackage{cmbright}')
rc('font', **{'family': 'sans-serif', 'sans-serif': ['Helvetica']})

%matplotlib inline

# This enables SVG graphics inline.
%config InlineBackend.figure_formats = {'png', 'retina'}

rc = {'lines.linewidth': 2,
      'axes.labelsize': 18,
      'axes.titlesize': 18,
      'axes.facecolor': 'DFDFE5'}
sns.set_context('notebook', rc=rc)
sns.set_style("dark")

mpl.rcParams['xtick.labelsize'] = 16
mpl.rcParams['ytick.labelsize'] = 16
mpl.rcParams['legend.fontsize'] = 14

sc.settings.verbosity = 3
gene_names = sc.queries.biomart_annotations('celegans', ['ensembl_gene_id',␣
 ↪'external_gene_name',
                                                                    'description',␣
 ↪'chromosome_name', 'gene_biotype'])
```

Load the data:

```
[2]: data = sc.read_loom('../data/All_Cells.loom')
data = data[~data.obs.Tissue.isin(['Unknown', 'Unannotated',␣
↪'Muscle_mesoderm'])]
data.var = data.var.join(gene_names.set_index('ensembl_gene_id'))
```

Next, we define the QC workflow. Moreover, the Taylor dataset is huge and enormously unbalanced (some tissues are much more over-represented over others). Therefore, I also wrote a subsampling function that will make the dataset smaller and also equalize the tissue representation across the dataset, to the extent that that is possible.

```
[3]: def qc(single_cell):
    """
    Performs a fairly standard QC workflow.
    Cells with <150 genes are filtered out.
    Genes expressed in <50 cells are removed.
    QC metrics for mitochondrial content, riboprotein content and NADH-enzymes␣
↪are calculated.
    Cells counts are normalized to counts per thousand (CPT). Because all cells␣
↪data ~300 counts / cell,
    CPTs are NOT log-transformed
    A subset scanpy object is returned.
    """
    data = single_cell.copy()
    sc.pp.filter_cells(data, min_genes=150)
    sc.pp.filter_genes(data, min_cells=50)
    data.var['mt'] = data.var.chromosome_name.fillna('').str.startswith('Mt') #␣
↪annotate the group of mitochondrial genes as 'mt'
    data.var['riboprots'] = data.var.external_gene_name.fillna('').str.
↪contains('^rp[sl]')
    data.var['nadh'] = data.var.description.fillna('').str.lower().str.
↪contains('nadh')
    sc.pp.calculate_qc_metrics(data, qc_vars=['mt', 'riboprots', 'nadh'],␣
↪percent_top=None, log1p=False, inplace=True)
    sc.pp.normalize_total(data, target_sum=10 ** 3)
    return data

def subsample(single_cell):
    """
    subsample all the cell types so that they are equally representeded.
    """
    index = []
    for tissue, df in single_cell.obs.groupby('Tissue'):
        print(tissue)
        if tissue in ['Rectal_cells', 'Excretory']:
            print('here')
```

```
            continue
        elif tissue is not 'Neuron':
            index += df.sample(np.min([df.shape[0], 2000])).index.tolist()
        else:
            index += df.sample(np.min([df.shape[0], 2000])).index.tolist()

    temp = single_cell[index].copy()
    sc.pp.highly_variable_genes(temp)
    sc.pp.scale(temp)
    sc.pp.pca(temp)
    return temp
```

In the next cell, I process the data using my `qc` function, and then I further subset cells to remove cells with <250 reads, that have too few or way too much mitochondrial, riboprotein or NADH-enzyme content. Neuronal cells are weird, so I remove neuronal cells with more than 800 counts per cell. Please note the Taylor *et al* dataset is unusual in that the average cell has VERY few counts, even for single-cell data.

[4]:

```
new = qc(data)

# subset data to keep only high quality stuff:
new = new[(new.obs.total_counts > 250) &
          (new.obs.pct_counts_mt.between(1, 10)) &
          (new.obs.pct_counts_riboprots.between(1, 20)) &
          (new.obs.pct_counts_nadh.between(.3, 3)) &
              ( ((new.obs.Tissue == 'Neuron') & (new.obs.total_counts < 800)) |
                (new.obs.Tissue != 'Neuron')
              )
         ]

sc.pl.violin(new, ['n_genes_by_counts', 'total_counts',],
             jitter=0.4, multi_panel=True)
sc.pl.violin(new, ['pct_counts_mt', 'pct_counts_nadh', 'pct_counts_riboprots'],
             jitter=0.4, multi_panel=True)
```

```
filtered out 19453 cells that have less than 150 genes expressed
filtered out 7018 genes that are detected in less than 50 cells
normalizing counts per cell
    finished (0:00:00)

/Users/davidangeles/opt/anaconda3/envs/scanpy/lib/python3.6/site-
packages/anndata/_core/anndata.py:1208: ImplicitModificationWarning:
Initializing view as actual.
  "Initializing view as actual.", ImplicitModificationWarning
Trying to set attribute `.obs` of view, copying.
… storing 'Cell.type' as categorical
Trying to set attribute `.obs` of view, copying.
… storing 'Detection' as categorical
Trying to set attribute `.obs` of view, copying.
```

```
… storing 'Experiment' as categorical
Trying to set attribute `.obs` of view, copying.
… storing 'Sample' as categorical
Trying to set attribute `.obs` of view, copying.
… storing 'Tissue' as categorical
Trying to set attribute `.obs` of view, copying.
… storing 'UMAP.clusters' as categorical
Trying to set attribute `.obs` of view, copying.
… storing 'birthtime' as categorical
Trying to set attribute `.obs` of view, copying.
… storing 'modality' as categorical
Trying to set attribute `.obs` of view, copying.
… storing 'neurotransmitter' as categorical
Trying to set attribute `.obs` of view, copying.
… storing 'partition' as categorical
Trying to set attribute `.var` of view, copying.
… storing 'external_gene_name' as categorical
Trying to set attribute `.var` of view, copying.
… storing 'description' as categorical
Trying to set attribute `.var` of view, copying.
… storing 'chromosome_name' as categorical
Trying to set attribute `.var` of view, copying.
… storing 'gene_biotype' as categorical
/Users/davidangeles/opt/anaconda3/envs/scanpy/lib/python3.6/site-
packages/seaborn/_core.py:1303: UserWarning: Vertical orientation ignored with
only `x` specified.
  warnings.warn(single_var_warning.format("Vertical", "x"))
/Users/davidangeles/opt/anaconda3/envs/scanpy/lib/python3.6/site-
packages/seaborn/_core.py:1303: UserWarning: Vertical orientation ignored with
only `x` specified.
  warnings.warn(single_var_warning.format("Vertical", "x"))
/Users/davidangeles/opt/anaconda3/envs/scanpy/lib/python3.6/site-
packages/seaborn/_core.py:1303: UserWarning: Vertical orientation ignored with
only `x` specified.
  warnings.warn(single_var_warning.format("Vertical", "x"))
/Users/davidangeles/opt/anaconda3/envs/scanpy/lib/python3.6/site-
packages/seaborn/_core.py:1303: UserWarning: Vertical orientation ignored with
only `x` specified.
  warnings.warn(single_var_warning.format("Vertical", "x"))
```

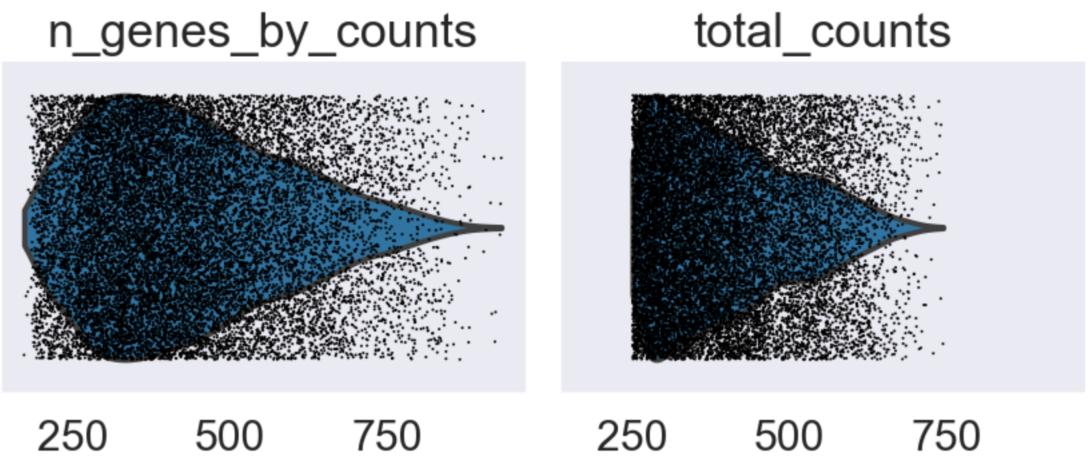


```
/Users/davidangeles/opt/anaconda3/envs/scanpy/lib/python3.6/site-
packages/seaborn/_core.py:1303: UserWarning: Vertical orientation ignored with
only `x` specified.
  warnings.warn(single_var_warning.format("Vertical", "x"))
/Users/davidangeles/opt/anaconda3/envs/scanpy/lib/python3.6/site-
packages/seaborn/_core.py:1303: UserWarning: Vertical orientation ignored with
only `x` specified.
  warnings.warn(single_var_warning.format("Vertical", "x"))
/Users/davidangeles/opt/anaconda3/envs/scanpy/lib/python3.6/site-
packages/seaborn/_core.py:1303: UserWarning: Vertical orientation ignored with
only `x` specified.
  warnings.warn(single_var_warning.format("Vertical", "x"))
/Users/davidangeles/opt/anaconda3/envs/scanpy/lib/python3.6/site-
packages/seaborn/_core.py:1303: UserWarning: Vertical orientation ignored with
only `x` specified.
  warnings.warn(single_var_warning.format("Vertical", "x"))
/Users/davidangeles/opt/anaconda3/envs/scanpy/lib/python3.6/site-
packages/seaborn/_core.py:1303: UserWarning: Vertical orientation ignored with
only `x` specified.
  warnings.warn(single_var_warning.format("Vertical", "x"))
/Users/davidangeles/opt/anaconda3/envs/scanpy/lib/python3.6/site-
packages/seaborn/_core.py:1303: UserWarning: Vertical orientation ignored with
only `x` specified.
  warnings.warn(single_var_warning.format("Vertical", "x"))
```

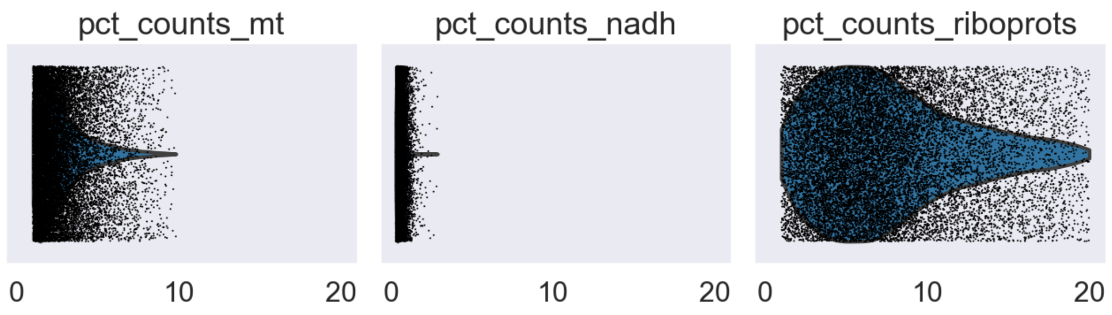


#### 1.1 QC Plots for single-cell rna-seq.

Let's make sure the cells look broadly OK:

[5]:
```
sc.pl.scatter(new, x='total_counts', y='n_genes_by_counts')
sc.pl.scatter(new, x='pct_counts_mt', y='n_genes_by_counts')
sc.pl.scatter(new, x='pct_counts_riboprots', y='n_genes_by_counts')
sc.pl.scatter(new, x='pct_counts_nadh', y='n_genes_by_counts')
sc.pl.scatter(new, x='pct_counts_mt', y='total_counts')
sc.pl.scatter(new, x='pct_counts_mt', y='pct_counts_nadh')
sc.pl.scatter(new, x='pct_counts_nadh', y='pct_counts_riboprots')
```

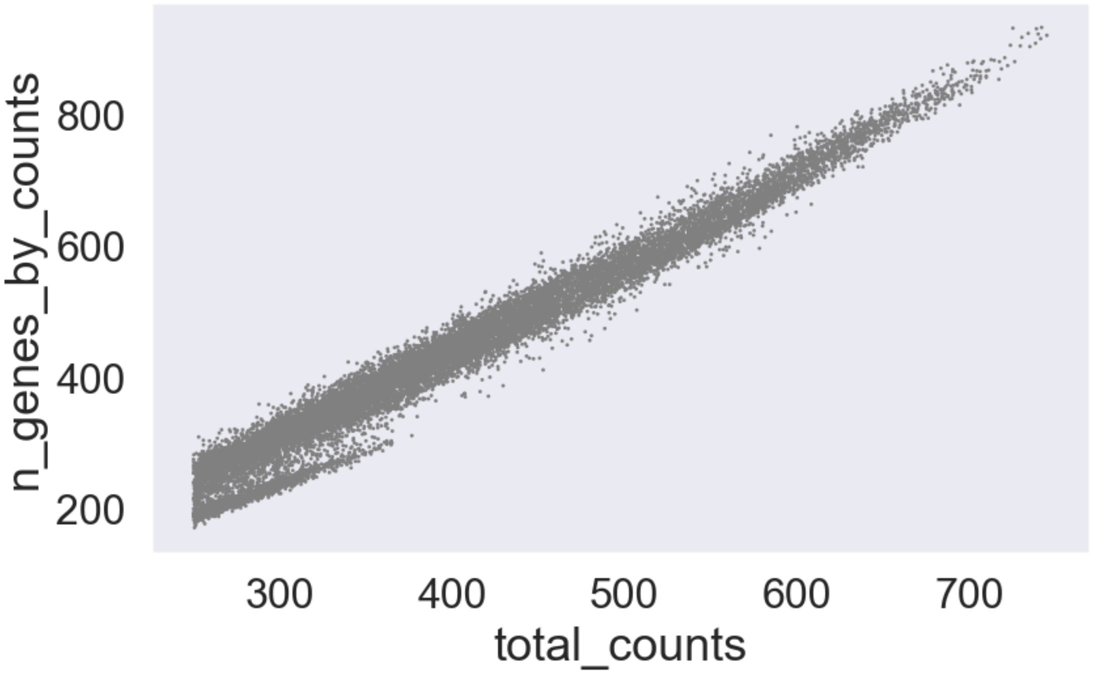

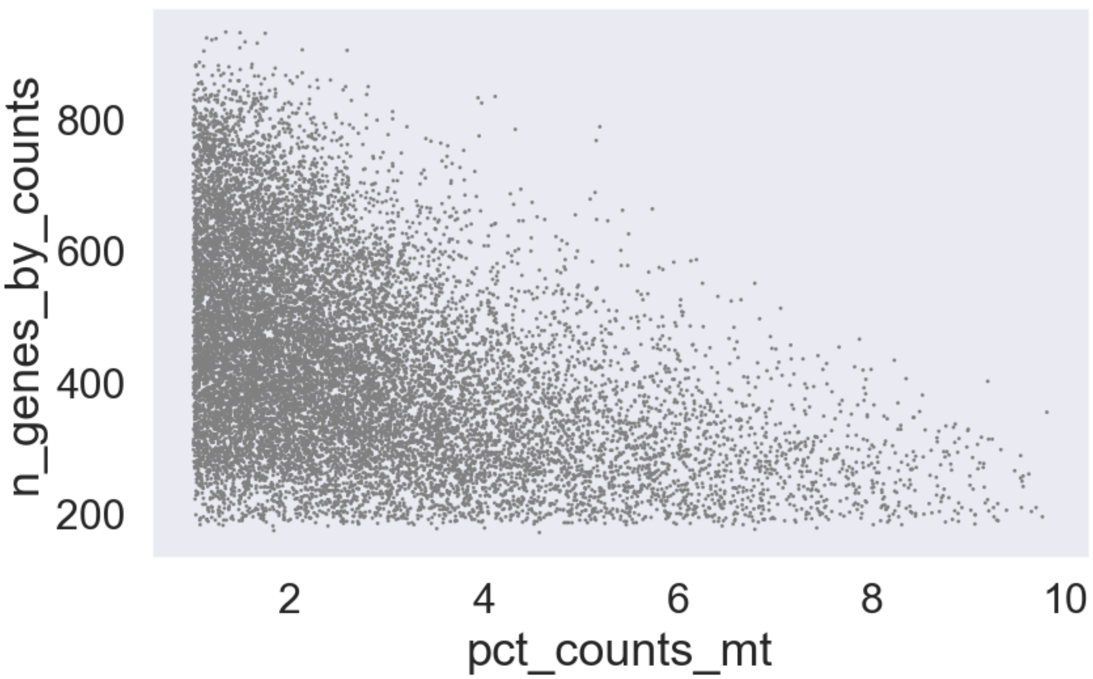
n_genes_by_counts
800
600
400
200
2
4
6
8
10
pct_counts_mt

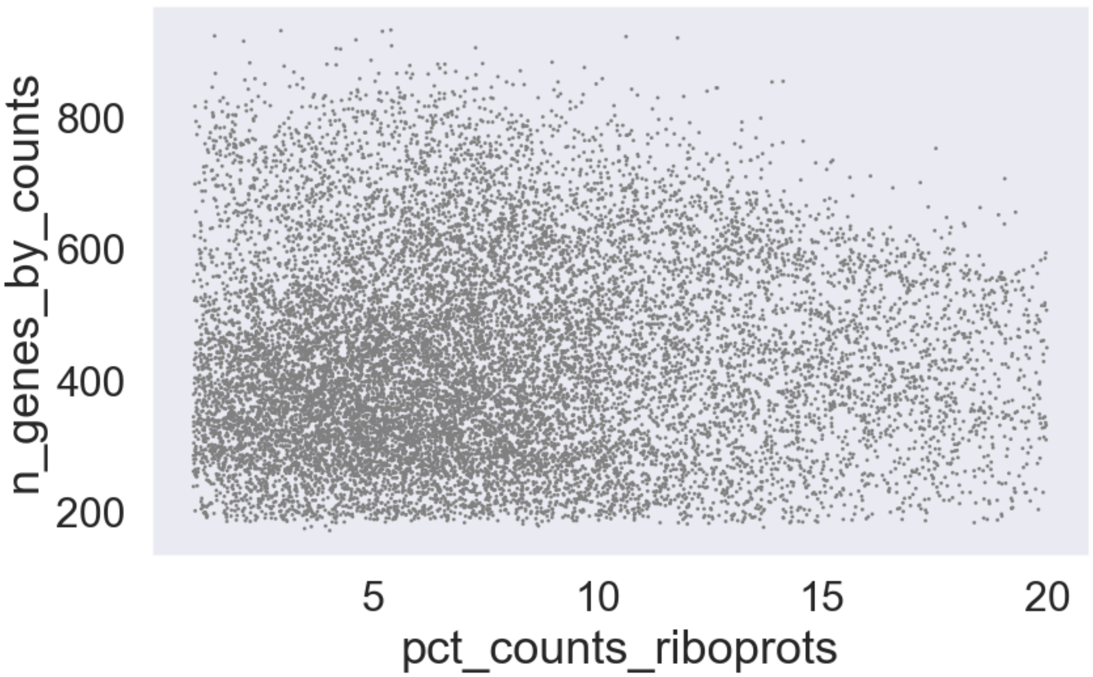
n_genes_by_counts
800
600
400
200
5
10
15
20
pct_counts_riboprots

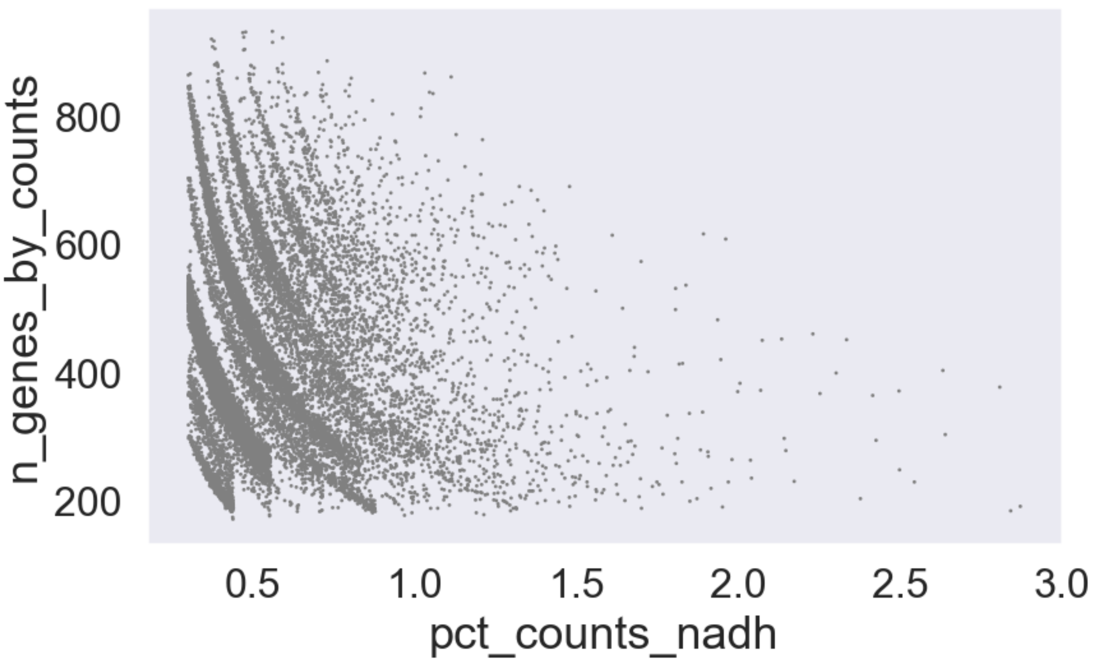
n_genes_by_counts
800
600
400
200
0.5
1.0
1.5
2.0
2.5
3.0
pct_counts_nadh

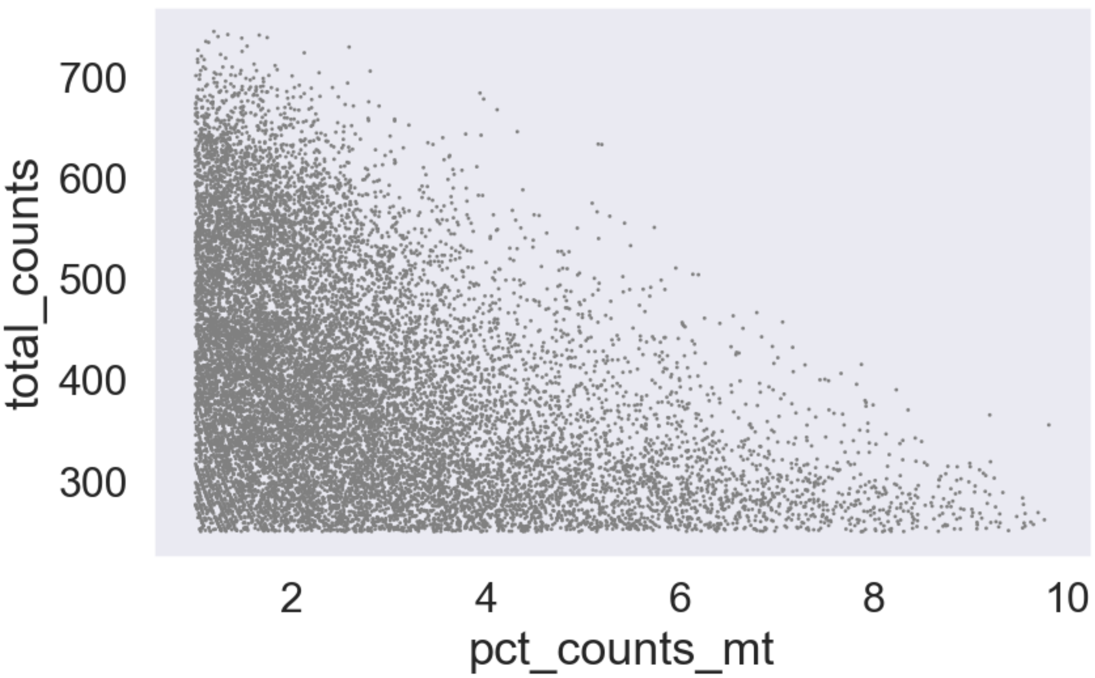
total_counts
700
600
500
400
300
2
4
6
8
10
pct_counts_mt

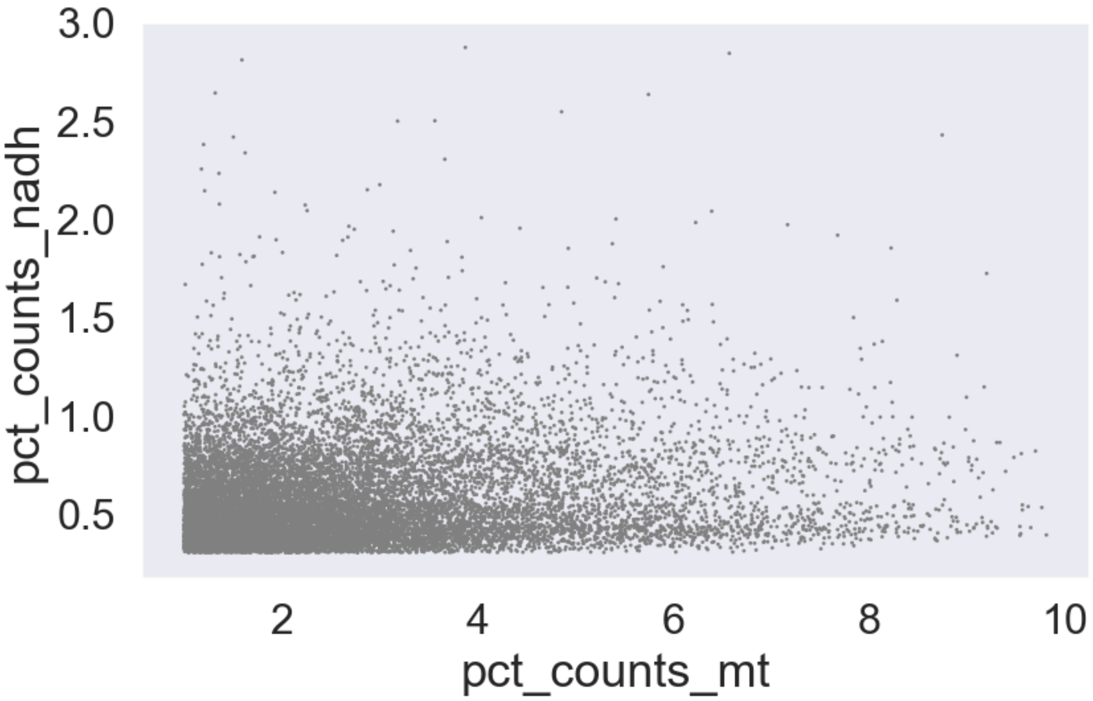

pct_counts_nadh
3.0
2.5
2.0
1.5
1.0
0.5
2
4
6
8
10
pct_counts_mt


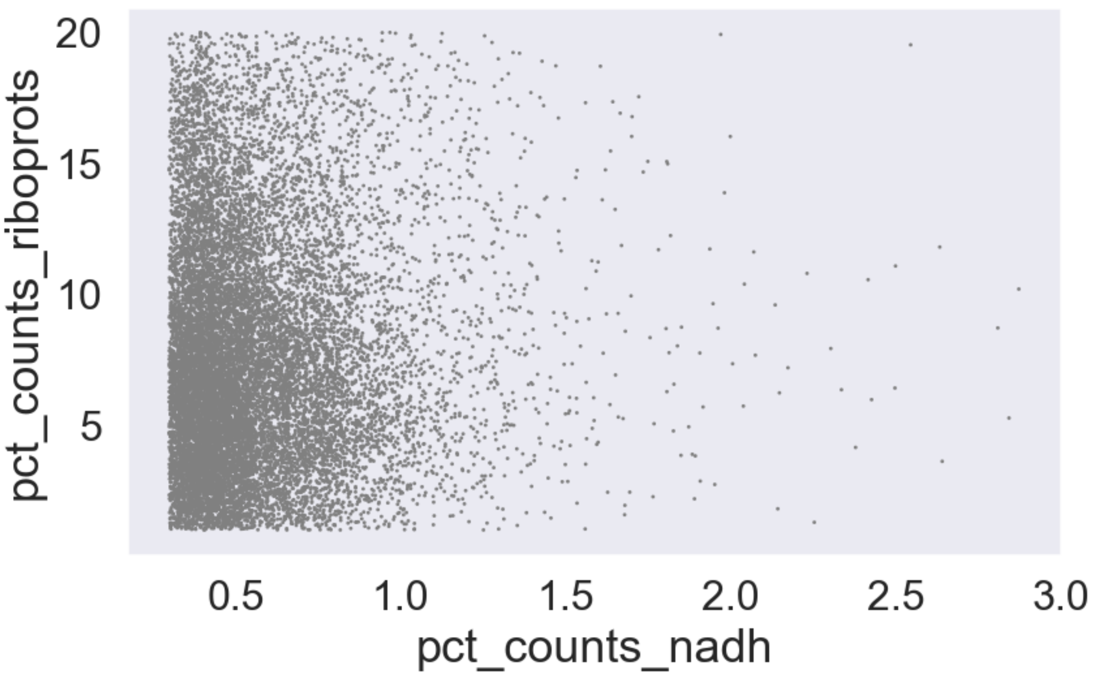

pct_counts_riboprots
20
15
10
5
0.5
1.0
1.5
2.0
2.5
3.0
pct_counts_nadh

#### 1.2 Subsample and embed single cell data

The workflow below is fairly self explanatory. PCA, clustering, UMAP to make sure nothing is glaringly wrong.

[6]:
```
new = subsample(new)
sc.pp.pca(new, n_comps=15)
sc.pp.neighbors(new, n_neighbors=100)
sc.tl.umap(new, min_dist=.2)
sc.pl.umap(new, color=['Tissue', 'total_counts'], wspace=0.4, alpha=0.2)
```

```
Excretory
here
Glia
Hypodermis
Intestine
Neuron
Pharynx
Rectal_cells
here
Reproductive
extracting highly variable genes
    finished (0:00:00)
--> added
    'highly_variable', boolean vector (adata.var)
    'means', float vector (adata.var)
    'dispersions', float vector (adata.var)
    'dispersions_norm', float vector (adata.var)
… as `zero_center=True`, sparse input is densified and may lead to large
memory consumption
computing PCA
    on highly variable genes
    with n_comps=50
    finished (0:00:01)
computing PCA
    on highly variable genes
    with n_comps=15
    finished (0:00:00)
computing neighbors
    using 'X_pca' with n_pcs = 15
    finished: added to `.uns['neighbors']`
    `.obsp['distances']`, distances for each pair of neighbors
    `.obsp['connectivities']`, weighted adjacency matrix (0:00:03)
computing UMAP
    finished: added
    'X_umap', UMAP coordinates (adata.obsm) (0:00:08)
```

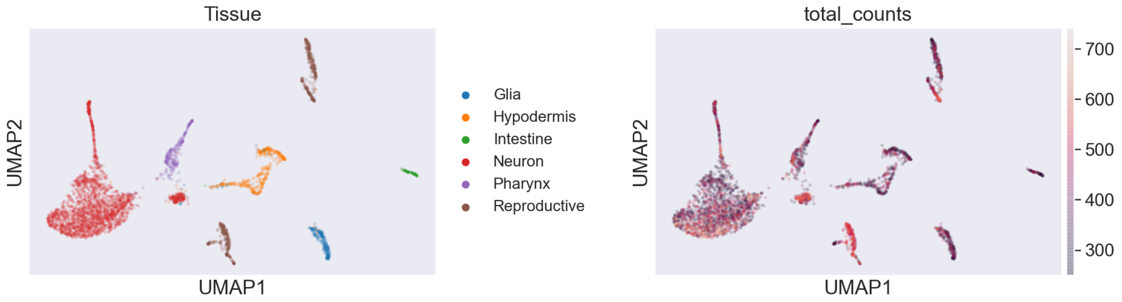


#### 1.3 Reformatting the labels and adjusting UMAP.

I want the sperm and oocyte/oocyte-precursor cells to be labeled independently. In the above, they are all pooled as 'reproductive'. I wrote a short function to split them up.

Next, we use PAGA to help guide the UMAP into a slightly better conformation, and we plot the new UMAP, colored by the fixed labels.

[7]:
```
def map_germline(x):
    """
    A way to rename the tissue samples such that the `Reproductive` term is␣
↪split into Germline and Sperm.
    """
    if x['Cell.type'] == 'Sperm':
        return 'Sperm'
    elif x['Cell.type'] == 'Germline':
        return 'Germline'
    elif x.Tissue == 'Reproductive':
        return 'Somatic Reproductive'
    else:
        return x.Tissue

new.obs['NewTissue'] = new.obs.apply(map_germline, axis=1)
```

[8]:
```
sc.tl.paga(new, groups='NewTissue')
sc.pl.paga(new)
sc.tl.umap(new, init_pos='paga', min_dist=1, spread=2)

sc.pl.umap(new, color=['NewTissue'])
```

```
… storing 'NewTissue' as categorical

running PAGA
    finished: added
    'paga/connectivities', connectivities adjacency (adata.uns)
    'paga/connectivities_tree', connectivities subtree (adata.uns) (0:00:00)
--> added 'pos', the PAGA positions (adata.uns['paga'])
```

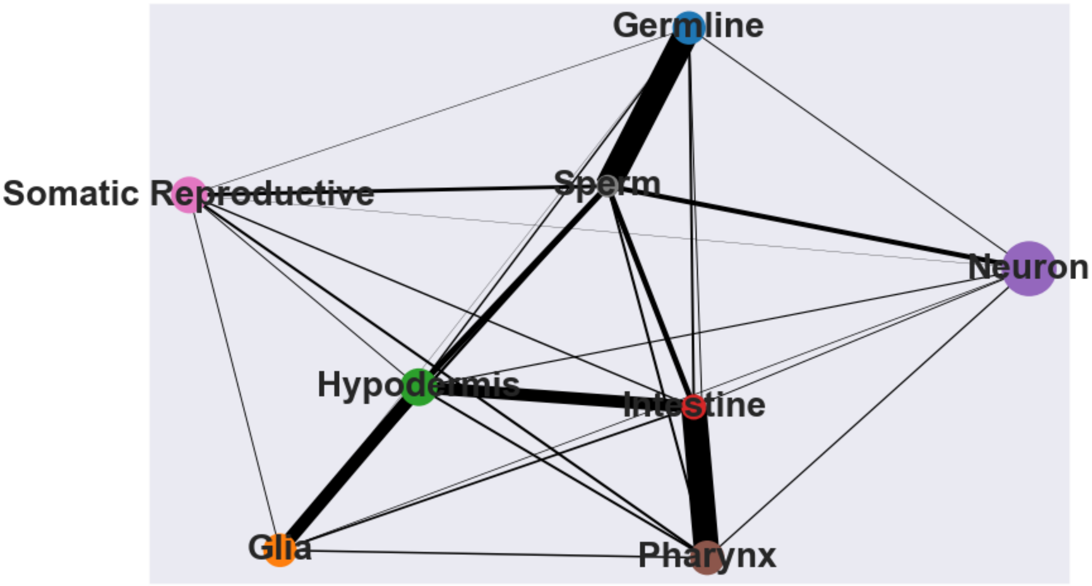


```
computing UMAP
    finished: added
    'X_umap', UMAP coordinates (adata.obsm) (0:00:07)
```

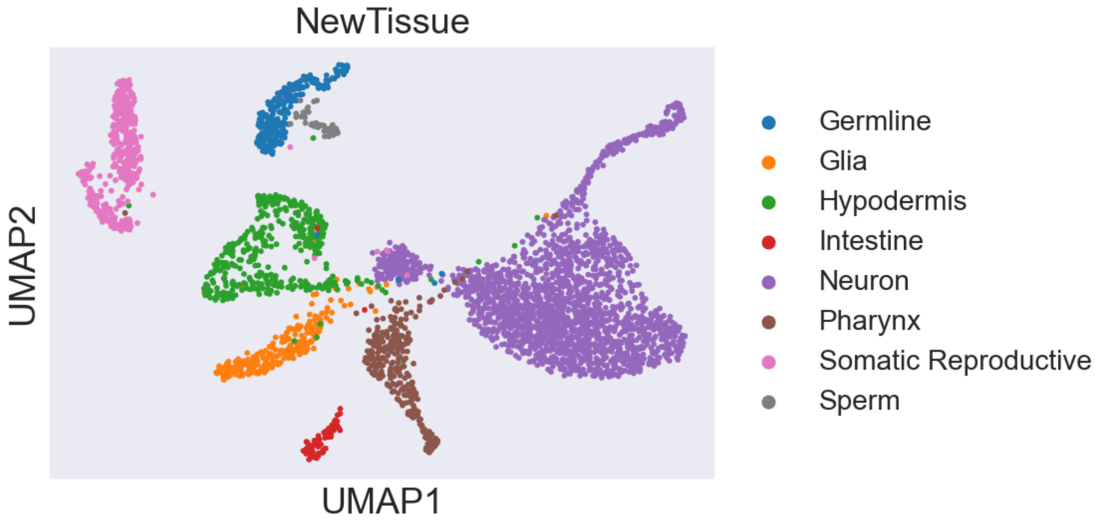


#### 1.4 Positive and negative ascr#10 signatures.

Next, we load our gene data and we set stringent conditions to find those genes that are most significantly up- or down-regulated in response to ascr#10 at either 50 or 58hrs. To be sure that whatever results we are looking at mean *something*, we also generated a random signature from random genes in our dataset.

[9]:
```
res = pd.read_csv('../data/master_table.tsv', sep='\t', index_col=0)
# keep DE genes only:
sel = (res['Sign-WT'] == 'Same') &\
      (res['padj-50'] < 0.01) & (res['padj-58'] < 0.01)
```

[10]:
```
pos_signature = res[sel & (res['log2FoldChange-58'] > 0)].index.values
neg_signature = res[sel & (res['log2FoldChange-58'] < 0)].index.values
random = res.sample(np.max([len(pos_signature), len(neg_signature)]),␣
↪replace=False).index.values
print(len(pos_signature), len(neg_signature))
```

```
200 148
```

[11]:
```
# print('positive genes found: ', len([p for p in pos_signature if p in new.
↪var_names]))
# print('negative genes found: ', len([p for p in neg_signature if p in new.
↪var_names]))
# print('random genes found: ', len([p for p in random if p in new.var_names]))

sel = (res['Sign-WT'] == 'Same') &\
      (res['padj-50'] < 0.1) & (res['padj-58'] < 0.1)

res[sel & (res['log2FoldChange-58'] > 0)].reset_index()['index'].to_csv('../
↪data/positive_signature.csv', index=False)
res[sel & (res['log2FoldChange-58'] < 0)].reset_index()['index'].to_csv('../
↪data/negative_signature.csv', index=False)
res.reset_index()['index'].to_csv('../data/background.csv', index=False)
```

I use the `score_genes` function in scanpy to generate a composite score for each of the positive, negative and random signatures we generated. Higher composite scores in a given tissue mean that the set of genes that participate in that score are expressed at higher levels relative to the bulk average expression. Thus, higher scores, in a sense, indicate some sort of tissue-specificity for our purposes.

[12]:
```
sc.tl.score_genes(new, pos_signature, score_name='PosAscrScore')
sc.tl.score_genes(new, neg_signature, score_name='NegAscrScore')
sc.tl.score_genes(new, random, score_name='RandomScore')
```

```
computing score 'PosAscrScore'
WARNING: genes are not in var_names and ignored: ['WBGene00003877',
'WBGene00010492']
    finished: added
    'PosAscrScore', score of gene set (adata.obs).
    1180 total control genes are used. (0:00:00)
computing score 'NegAscrScore'
WARNING: genes are not in var_names and ignored: ['WBGene00000169',
'WBGene00000830', 'WBGene00000831', 'WBGene00002174', 'WBGene00004904',
```

```
'WBGene00006464', 'WBGene00007097', 'WBGene00007228', 'WBGene00007336',
'WBGene00008425', 'WBGene00008499', 'WBGene00008724', 'WBGene00010241',
'WBGene00010366', 'WBGene00010540', 'WBGene00011821', 'WBGene00011910',
'WBGene00012013', 'WBGene00012531', 'WBGene00012809', 'WBGene00013771',
'WBGene00013996', 'WBGene00014179', 'WBGene00014665', 'WBGene00016174',
'WBGene00017071', 'WBGene00017364', 'WBGene00018134', 'WBGene00019086',
'WBGene00020223', 'WBGene00020353', 'WBGene00020414', 'WBGene00021007',
'WBGene00021167', 'WBGene00021381', 'WBGene00021969', 'WBGene00022090',
'WBGene00045338', 'WBGene00219376']
    finished: added
    'NegAscrScore', score of gene set (adata.obs).
    1090 total control genes are used. (0:00:00)
computing score 'RandomScore'
WARNING: genes are not in var_names and ignored: ['WBGene00016763',
'WBGene00044512', 'WBGene00004904', 'WBGene00011190', 'WBGene00017667',
'WBGene00016967', 'WBGene00009182', 'WBGene00003943', 'WBGene00019970',
'WBGene00001795', 'WBGene00015138', 'WBGene00303075', 'WBGene00012151',
'WBGene00010773', 'WBGene00269384', 'WBGene00004242', 'WBGene00007139',
'WBGene00005228', 'WBGene00001514', 'WBGene00007598', 'WBGene00010540',
'WBGene00020879', 'WBGene00012164', 'WBGene00016758', 'WBGene00009447',
'WBGene00002174', 'WBGene00044337', 'WBGene00012103', 'WBGene00021713',
'WBGene00014029', 'WBGene00017054', 'WBGene00009548', 'WBGene00008350',
'WBGene00185002', 'WBGene00001511', 'WBGene00019341', 'WBGene00011801',
'WBGene00007789', 'WBGene00014179', 'WBGene00021354', 'WBGene00016823',
'WBGene00018342', 'WBGene00017048', 'WBGene00021878', 'WBGene00008917',
'WBGene00044032', 'WBGene00010516']
    finished: added
    'RandomScore', score of gene set (adata.obs).
    1130 total control genes are used. (0:00:00)
```

[13]:
```
sns.boxplot(y='Tissue', x='PosAscrScore', data=new.obs)
plt.xlim(-2, 2)
plt.axvline(0, color='red')
scores = new.obs.groupby('Tissue').PosAscrScore.apply(np.mean)
scores = scores / scores.max()
scores.sort_values()
```

[13]:
```
Tissue
Glia           -0.436641
Neuron         -0.265261
Hypodermis     -0.097456
Intestine       0.097343
Pharynx         0.124444
Reproductive    1.000000
Name: PosAscrScore, dtype: float64
```

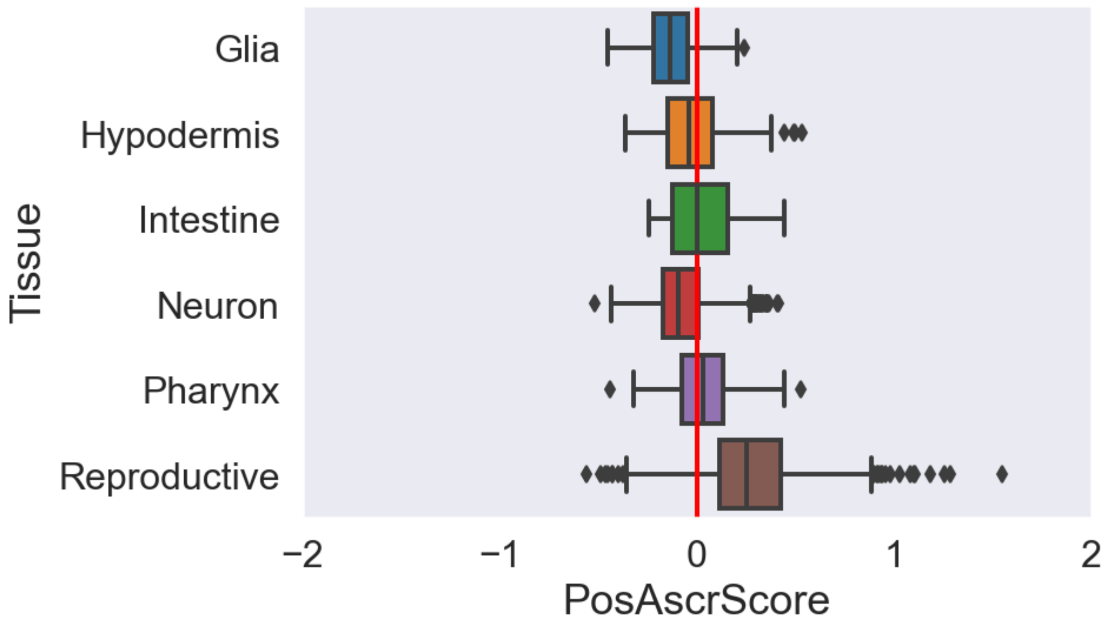


[14]:
```
sns.boxplot(y='Tissue', x='NegAscrScore', data=new.obs)
plt.xlim(-2, 2)

neg_scores = new.obs.groupby('Tissue').NegAscrScore.apply(np.mean)
neg_scores = neg_scores / neg_scores.max()
neg_scores.sort_values()
```

[14]:
```
Tissue
Pharynx        -0.135059
Neuron         -0.128662
Reproductive    0.022506
Glia            0.079274
Hypodermis      0.471763
Intestine       1.000000
Name: NegAscrScore, dtype: float64
```

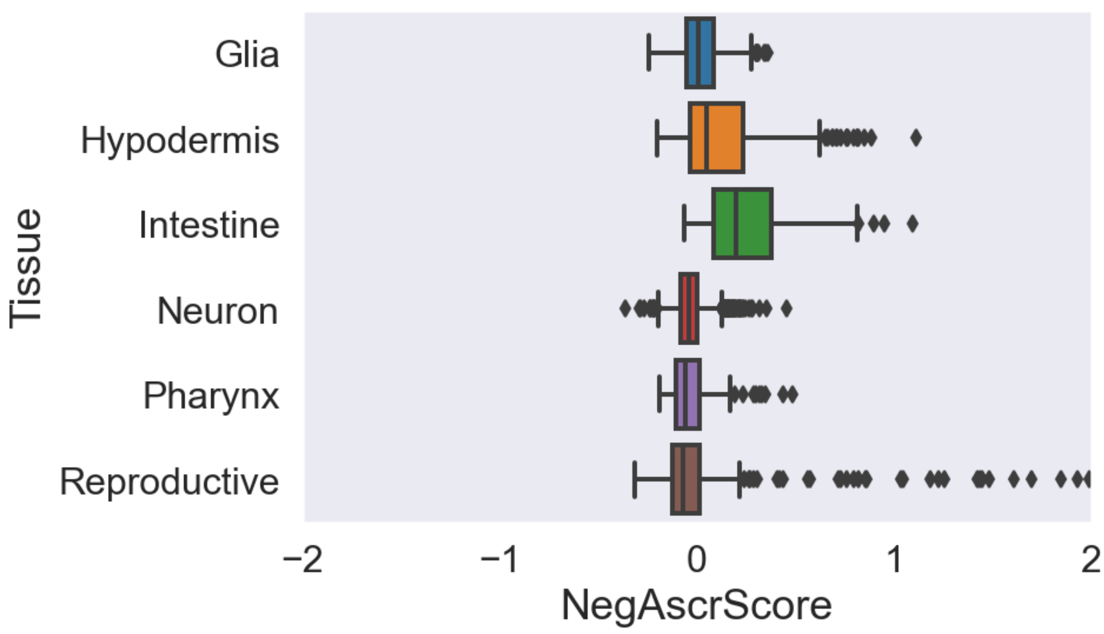


[15]:
```
sns.boxplot(y='Tissue', x='RandomScore', data=new.obs)
plt.xlim(-2, 2)
```

[15]: (-2.0, 2.0)

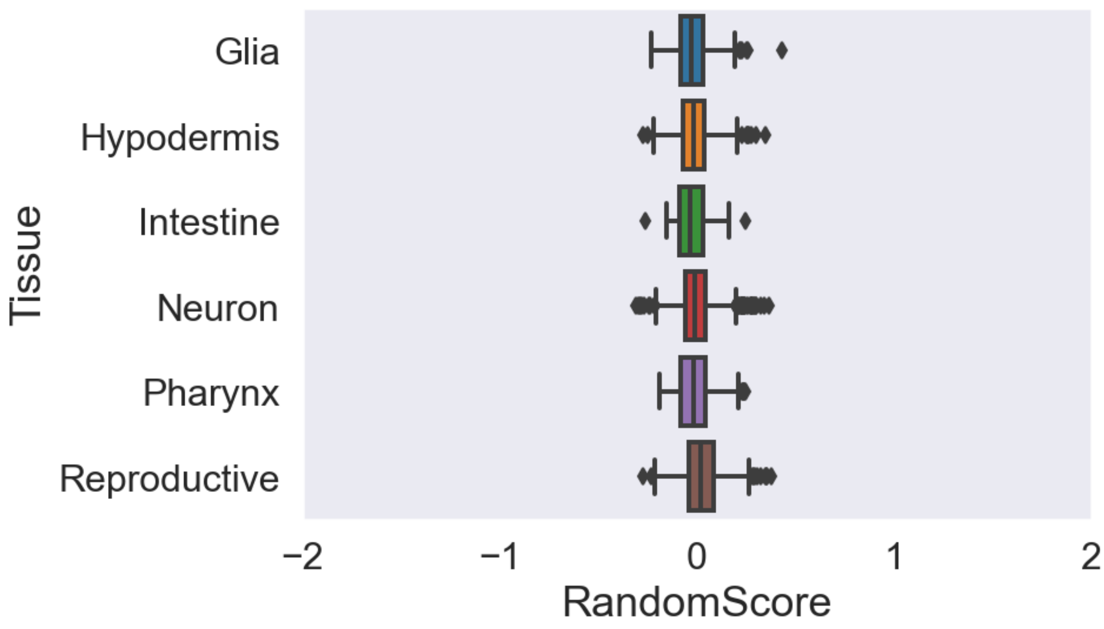

Let's put it all together to generate the colored UMAPs for our paper figures:

[16]:
```
sc.pl.umap(new, color=['NewTissue', 'PosAscrScore'], cmap='viridis', vmax=0.5,␣
↪vmin=0, wspace=.35, save='_panel_1.svg')
sc.pl.umap(new, color=['NegAscrScore', 'RandomScore'], cmap='viridis', vmax=.5,␣
↪vmin=0, wspace=.35, save='panel_2.svg')
```

```
WARNING: saving figure to file figures/umap_panel_1.svg
```

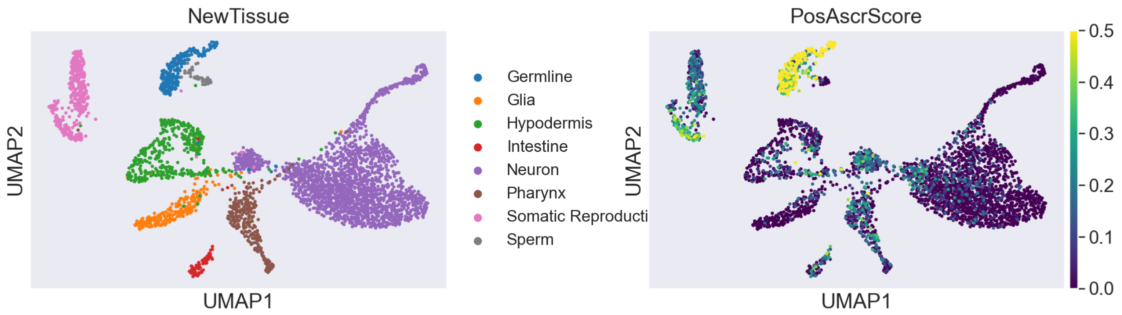


```
WARNING: saving figure to file figures/umappanel_2.svg
```

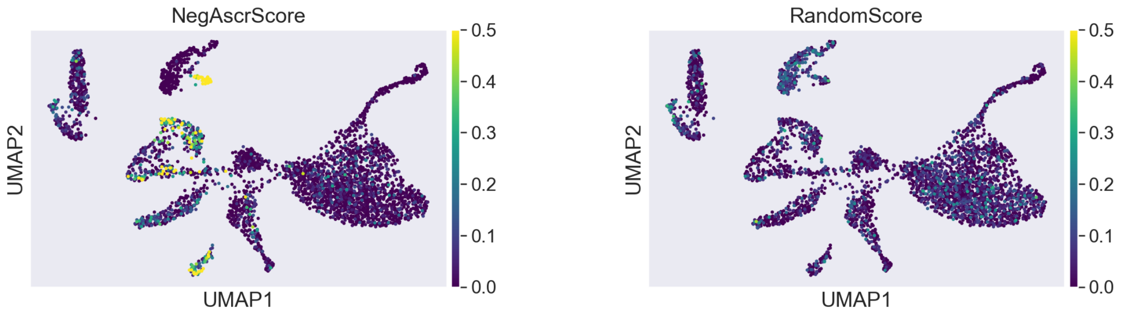


Finally, just to make absolutely sure our interpretations are correct, let's make a density UMAP plot to reveal where the germline / sperm cells are located in the UMAP plot.

[17]:
```
def map_germline(x):
    """A way to drop all labels that are not sperm or germline."""
    if x in ['Sperm', 'Germline']:
        return x
    else:
        return ''

new.obs['Germ'] = new.obs['Cell.type'].apply(map_germline)
```

[18]:
```
# reality check to look where the sexual cells are going in the UMAP.
sc.tl.embedding_density(new, basis='umap', groupby='Germ')
sc.pl.embedding_density(
    new, basis='umap', key='umap_density_Germ', group=['Sperm', 'Germline']
)
```

```
… storing 'Germ' as categorical

computing density on 'umap'
--> added
    'umap_density_Germ', densities (adata.obs)
    'umap_density_Germ_params', parameter (adata.uns)

/Users/davidangeles/opt/anaconda3/envs/scanpy/lib/python3.6/site-
packages/scanpy/plotting/_tools/__init__.py:1156: MatplotlibDeprecationWarning:
You are modifying the state of a globally registered colormap. In future
versions, you will not be able to modify a registered colormap in-place. To
remove this warning, you can make a copy of the colormap first. cmap =
copy.copy(mpl.cm.get_cmap("YlOrRd"))
  color_map.set_over('black')
/Users/davidangeles/opt/anaconda3/envs/scanpy/lib/python3.6/site-
packages/scanpy/plotting/_tools/__init__.py:1157: MatplotlibDeprecationWarning:
You are modifying the state of a globally registered colormap. In future
versions, you will not be able to modify a registered colormap in-place. To
remove this warning, you can make a copy of the colormap first. cmap =
copy.copy(mpl.cm.get_cmap("YlOrRd"))
  color_map.set_under('lightgray')
/Users/davidangeles/opt/anaconda3/envs/scanpy/lib/python3.6/site-
packages/scanpy/plotting/_tools/scatterplots.py:400:
MatplotlibDeprecationWarning: Passing parameters norm and vmin/vmax
simultaneously is deprecated since 3.3 and will become an error two minor
releases later. Please pass vmin/vmax directly to the norm when creating it.
  **kwargs,
```

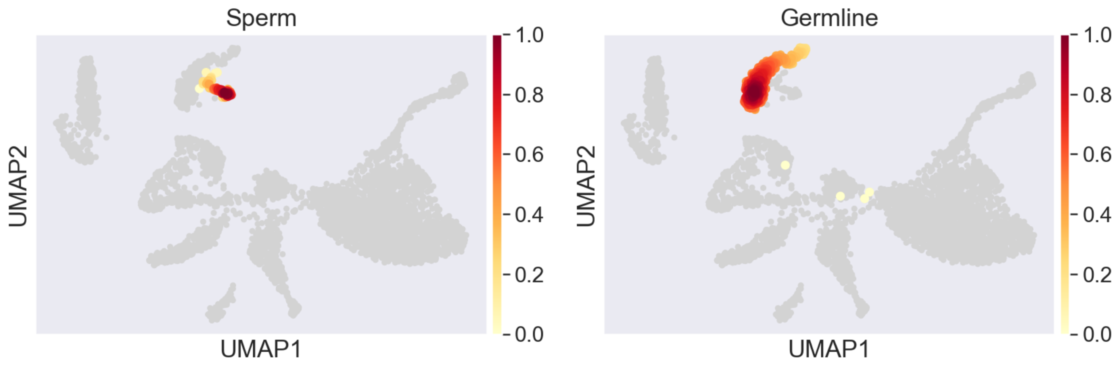


It appears that the sperm and oocyte cells are actually clustered where they appear to be. It's not an artifact! Yay!
